# Integrated multi-omics profiling reveals the role of the DNA methylation landscape in shaping biological heterogeneity and clinical behaviour of metastatic melanoma

**DOI:** 10.1101/2025.02.07.637045

**Authors:** Andrea Anichini, Francesca P. Caruso, Vincenzo Lagano, Teresa M.R. Noviello, Rossella Tufano, Gabriella Nicolini, Alessandra Molla, Ilaria Bersani, Francesco Sgambelluri, Alessia Covre, Maria F. Lofiego, Sandra Coral, Anna Maria Di Giacomo, Elena Simonetti, Barbara Valeri, Mara Cossa, Filippo Ugolini, Sara Simi, Daniela Massi, Massimo Milione, Andrea Maurichi, Roberto Patuzzo, Mario Santinami, Michele Maio, Michele Ceccarelli, Roberta Mortarini, the EPigenetic Immune-oncology Consortium Airc (EPICA) investigators

## Abstract

The biological and clinical relevance of the DNA methylation landscape in metastatic melanoma (MM) remains underexplored. In a retrospective cohort of 191 MM lesions from 165 AJCC Stage III and IV patients (EPICA cohort) we identified four tumor subsets (i.e. DEMethylated, LOW, INTermediate and CIMP) with progressively increasing levels of DNA methylation. These findings were validated in the TCGA MM. In EPICA, patients with LOW methylation tumors exhibited a significantly longer survival and a lower progression rate to more advanced AJCC stages, compared to patients with CIMP tumors. Furthermore, in an independent adjuvant immune checkpoint blockade MM cohort, patients with DEM/LOW pre-therapy lesions showed significantly longer relapse-free survival compared to those with INT/CIMP lesions. RNA-seq data analysis revealed that LOW and CIMP EPICA tumors showed opposite activation of master molecules influencing prognostic target genes, and differential expression of immunotherapy response and melanoma differentiation signatures. Compared to CIMP tumors, LOW lesions showed enrichment for pre-exhausted and exhausted T cells and more frequent retention of HLA Class I antigens. The differentiation and immune-related transcriptional features associated with LOW vs CIMP lesions were tumor-intrinsic programs retained in-vitro by melanoma cell lines. Consistently, treatment of differentiated melanoma cell lines with a DNMT inhibitor induced global DNA de-methylation, promoted de-differentiation and upregulated viral mimicry and IFNG predictive signatures of immunotherapy response. These findings underscore the role of DNA methylation in driving MM biological and clinical heterogeneity and support exploration of methylome targeting strategies for precision immunotherapy in melanoma.

## Introduction

Nonmutational epigenetic reprogramming, one of the enabling characteristics contributing to the acquisition of the hallmarks of cancer^1^, refers to the plethora of epigenetic mechanisms affecting gene expression independently from genome instability and gene mutation. Among such mechanisms, altered DNA methylation at CpG dinucleotides contributes to tumorigenesis and to cancer evolution^2^. Early studies identified a progressive reduction in the overall 5-methylcytosine content along with the transition from benign lesions to primary malignant tumors and then to metastatic lesions^3^. Conversely, cancer-specific DNA hypermethylation at CpG islands, associated with transcriptional silencing of tumor suppressors, was subsequently discovered^4^. In cutaneous melanoma, both global and gene-specific DNA methylation changes play a role in melanocyte transformation^5^, contribute to shape the melanoma differentiation phenotype^6,7^, impact tumor progression^8–10^ and associate with clinical outcome^8,11–15^.

DNA methylation profiling across melanoma cohorts can also classify lesions into distinct subgroups. Analysis of two primary melanomas cohorts (n=47 and n=89) by Thomas et al.^16^ and Conway et al.^14^ revealed three methylation classes (Low, Intermediate, High/CIMP). In the first cohort^16^ the highly methylated class was positively associated with Breslow thickness. In the second cohort^14^ the CIMP class was associated with higher AJCC Stage, age at diagnosis ≥65 years, lower tumor-infiltrating lymphocyte grade and worse melanoma-specific survival. More recently, a two-fold higher likelihood of 5-year death for patients with primary melanomas in the CIMP or intermediate methylation classes, compared to the Low methylation class, has been reported in a cohort of 422 stage II and III patients^17^. In n=50 metastatic melanomas (MM) Lauss et al.^15^ identified three methylation groups (MS1-3). MS1 and MS3 showed respectively the highest and the lowest global methylation levels associated with differential expression of immune-related and proliferative signatures^15^. The Cancer Genome ATLAS (TCGA) network^18^ identified four methylation subsets, with increasing global methylation levels (hypomethylated, normal-like, hypermethylated and CIMP) in a cohort of n=332 primary and MM lesions from previously untreated patients. The normal-like subset was associated with a longer survival compared to the CIMP class, although patients were not stratified for tumor stage^18^.

Despite these initial intriguing insights, the overall biological significance and potential therapeutic relevance of methylation-defined subsets in melanoma remain underexplored, as several key gaps in knowledge still need to be addressed. First, the impact of DNA methylation classes on the clinical outcome of patients, and on disease progression to more advanced stages, has poorly understood molecular underpinnings. Second, the potential association of the methylation subsets with transcriptional programs of response to immune checkpoint blockade (ICB) therapies and with the structure of the immune contexture remain to be fully deciphered. Third, the role of DNA methylation in identifying patients that represent optimal candidates for adjuvant ICB therapies is unknown. Fourth, the direct role of DNA methylation in shaping melanoma transcriptional programs of immunity and differentiation needs to be verified by cause-effect experiments. Fifth, the development of more effective immunotherapeutic approaches is urgently needed for advanced melanoma, therefore it is crucial to verify whether DNA methylation of melanoma is a suitable target for immune reprogramming of tumors. To address these issues, we carried out an integrative analysis of multi-omics data generated in tumor lesions from a fully annotated cohort of Stage III/IV patients (EPICA). Among the four methylation-defined subsets that were identified, the LOW and CIMP lesions expressed opposite melanoma differentiation and immune-related programs, distinct tumor immune contextures and divergent clinical evolution. Results from an independent adjuvant ICB MM cohort supported the clinical relevance of the DNA methylation classification. Moreover, global DNA demethylation, induced by a DNMT inhibitor (DNMTi), reprogrammed MITF^HI^/PMEL^HI^ melanoma cells towards a de-differentiated profile associated with upregulation of immune-related signatures.

## Results

### DNA methylation profiling of MM lesions reveals four subsets

We used reduced representation bisulfite sequencing (RRBS), RNA sequencing (RNA-seq) and whole exome sequencing (WES) on a fully annotated cohort of n=191 melanoma lesions from n=165 AJCC 8^th^ edition stage III and/or IV patients (thereafter “EPICA cohort”) who underwent surgery for the resection of metastatic lesions between 09/2002 and 12/2017 (Supplementary methods). Median follow-up was 28.4 months (range: 1.9 to 194.3 months from surgery for removal of the investigated lesions to death or to last follow-up) and 98.2% of the patients had not received any previous medical/systemic treatment (Supplementary Table 1a,b for all clinical and pathological data). By unbiased clustering of the top 1% most variable CpG sites (n=4,064, Supplementary Table 2) we identified four classes with progressive increase in global methylation levels: demethylated (DEM), LOW, intermediate (INT) and hypermethylated (CIMP) (Fig. 1a). In the whole EPICA cohort the mean tumor purity, assessed by digital pathology on H&E-stained sections, was 83.1% (Supplementary Table 1b) and the CIMP class showed slightly higher tumor purity values compared to the DEM and LOW subsets (Extended Data Fig. 1a). We identified four methylation classes even in the TCGA Firehose Legacy melanoma cohort (www.cbioportal.org) of 368 MM lesions (Extended Data Fig. 1b) and in the TCGA cohort of 104 primary melanoma lesions (Extended Data Fig. 1c). In the EPICA cohort 103 out of 165 patients (62.4%) underwent further clinical stage progression during follow-up (Fig. 1b and Supplementary Table 1a). However, among patients who did not progress to any subsequent AJCC stage, 38.03% had LOW lesions, compared to 19.7%, 25.4% and 16.9% of non-progressors among patients with DEM, INT or CIMP lesions, respectively (Fig. 1c). The proportion of patients progressing to CNS metastases (AJCC Stage IVM1D) in the DEM, INT and CIMP clusters was respectively 25.0%, 17.3% and 26.5%, compared to 9.3% in the LOW cluster (Supplementary Table 1b). In both the EPICA and the TCGA MM cohorts, patients in the LOW and CIMP subsets showed the longest and the shortest survival, respectively (Fig. 1d,e). Exome sequencing profiling of the EPICA cohort lesions revealed that mutations in the most frequently altered genes, such as BRAF and NRAS, were not associated with the methylation subsets (Extended Data Fig. 2). Only 6 genes (*RYR3, MGAM, SAMD9L, PAPPA2, TENM4, SNF208*) showed significant associations with the methylation classes. Moreover, the four methylation classes did not differ significantly in terms of the tumor mutational burden (TMB) value (Extended Data Fig. 2; p=0.3 by Anova).

**Figure 1.**
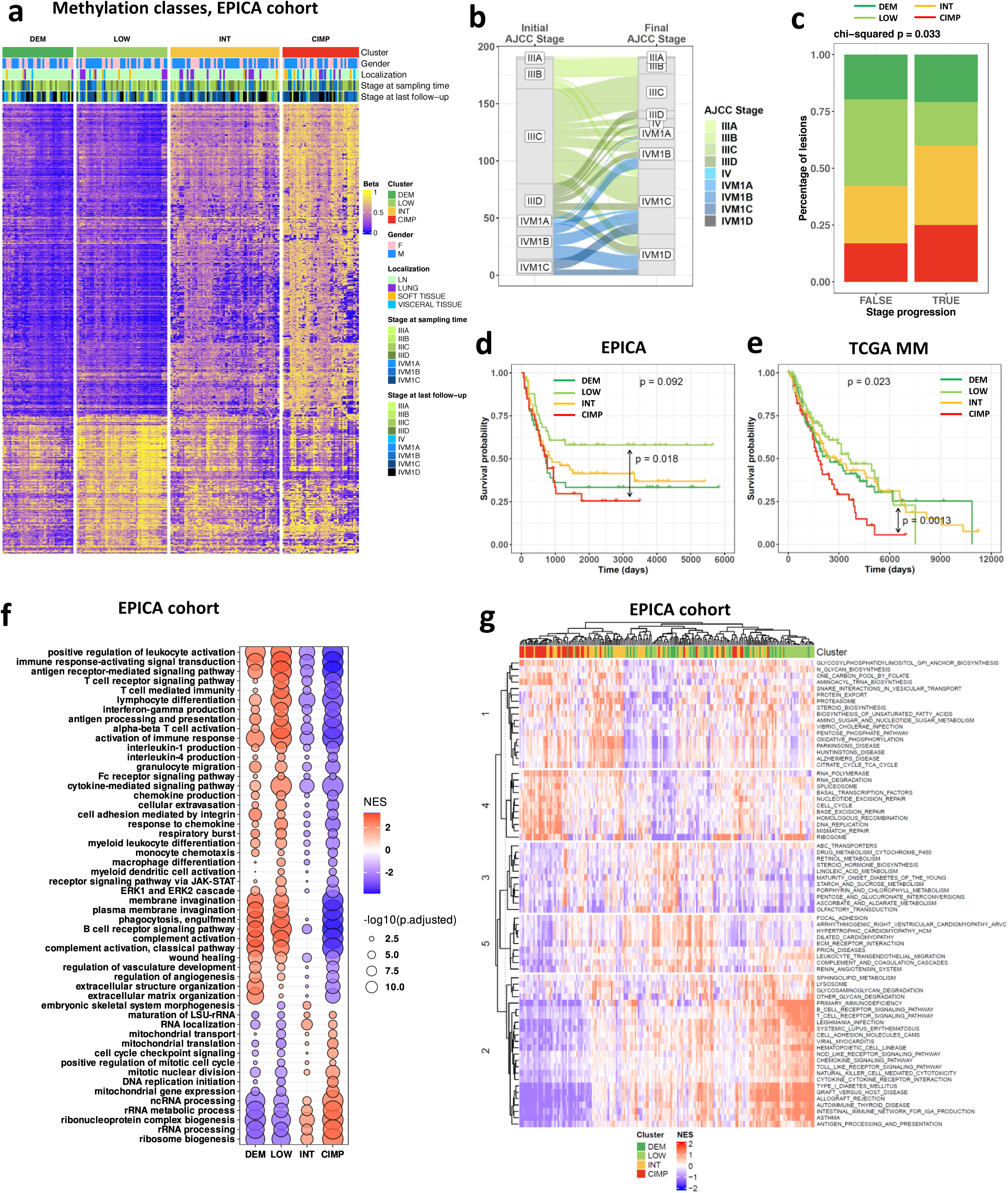
DNA Methylation profiling of MM EPICA cohort identifies four classes: clinical relevance and transcriptional programs. **a,** Consensus clustering of n=191 lesions from n=165 AJCC stage III/IV MM patients (EPICA cohort) based on 4064 most variable CpG sites. **b,** Alluvial plot showing the initial AJCC stage of n=165 patients in the EPICA cohort and the final AJCC stage at the end of follow-up. **c**, Stacked bar plot showing the percentage of samples in EPICA cohort according to stage progression occurring (TRUE) or not (FALSE) after the surgical resection of the initial lesion used for methylation profiling. **d**,**e**, Kaplan-Meier survival curves as a function of methylation cluster in EPICA cohort (d, n=165) and in TCGA MM cohort (e, n=368). Statistical analysis by Chi-square (c) and log rank test (d,e). **f,** Dot plot of the normalized enrichment score (NES) of significant Biological Processes (BP) obtained from GSEA (adjusted p-value < 0.05) in DEM, LOW, INT and CIMP methylation-defined classes of n=187 lesions from n=165 patients of the EPICA cohort. Dot color represent NES and the size represent p-value resulting from each comparison (one group vs all the others). GO terms selected are the top and bottom 25 significant for each comparison. **g,** Unsupervised clustering of 191 samples of the EPICA cohort based on 72 KEGG pathways significantly enriched in at least the 80% of samples (absolute logit NES >0.58 and p-value < 0.05).

### Transcriptional landscape of methylation subsets in metastatic melanoma

By integrating DNA methylation results with RNA-seq data in the EPICA cohort, we tested the relationship linking promoter methylation to gene expression in the four methylation clusters. By over-representation analysis we looked at distinct collections of gene sets (GO, HP, KEGG and REACT). In the LOW class, genes belonging to a large set of immune-related pathways were up-regulated and hypomethylated (Extended Data Fig. 3a), while genes belonging to developmental pathways were down-regulated and hypermethylated (Extended Data Fig. 3b). In contrast, genes belonging to cell-cell communication and developmental processes were up-regulated and hypomethylated in the DEM subset (Extended Data Fig. 3a). In the CIMP class we found enrichment for down-regulated and hypermethylated gene sets belonging to immune-related pathways (Extended Data Fig. 3b). Therefore, immune-related pathways showed opposite methylation/expression relationships in the LOW vs CIMP classes. In agreement with these findings, by GSEA we found that the DEM and the LOW subsets were enriched for immune-related biological processes, while the CIMP and INT subsets were characterized by mitochondrial metabolism and cell cycle/proliferation pathways (Fig. 1f). An unsupervised pathway-based deconvolution algorithm^19^ applied to 186 KEGG pathways, 72 of which were significantly enriched, segregated CIMP samples from LOW samples, identifying two extreme functional states confirming the association of cell cycle/proliferation pathways with CIMP and INT subsets and of immune-related pathways with LOW and DEM subsets (Fig. 1g). These findings support the notion that DNA methylation shapes distinct transcriptional phenotypes of the MM subsets.

### Immune-related master molecules with differential activation in DNA methylation classes control gene signatures with prognostic relevance

We used Ingenuity Pathway Analysis (IPA) as described^20^, on differentially expressed genes in the four methylation classes of EPICA (Supplementary Table 3). We found that the DEM and LOW subsets were characterized by activation of master molecules controlling innate and adaptive immunity (such as IFNG, TNF, IL1B), while the INT and CIMP subsets were characterized by inhibition of these regulators (Supplementary Fig. 1). By the IPA “Upstream Regulator” (UR) computational tool^20^ we identified master molecules explaining the differences in gene expression among methylation subsets. LOW and CIMP lesions showed an opposite pattern of UR activation vs UR inhibition (Fig. 2a). In the LOW class top activated UR (e.g., IFNG, TNF, STAT1, NFkB, IFNA, STING1) were positive regulators of innate and adaptive immunity (Fig. 2a). In contrast, several top activated UR in CIMP subset (e.g., TREX1, IRGM, IL1RN, RNASEH2B, HIVEP1, SIRT1), were negative regulators of innate immunity pathways^21–25^ (Fig. 2a). Activation of top UR as IFNG in LOW lesions reflected increased IFNG target gene expression in LOW compared to CIMP lesions (Extended Data Fig. 4). Conversely, due to their negative immune function, activation of top UR as TREX1, IRGM and IL1RN in CIMP lesions reflected reduced target gene expression in CIMP compared to LOW lesions (Extended Data Fig. 4). In agreement, the target gene signatures of these four UR were expressed above median Z score value of each signature in the majority of LOW lesions and below the median z score value in the majority of CIMP lesions (Fig. 2 b,c and Extended Data Fig. 5 a,c). By using the “clinical outcome” module of the TIMER 2.0 web application^26^, we found that the majority of the target genes of these four UR had prognostic significance in the TCGA MM dataset and conferred reduced risk (Extended Data Fig.4). In agreement, we found that the target gene signatures of these four UR had prognostic significance in both the EPICA and TCGA MM cohorts (Fig. 2d,e and Extended Data Fig. 5b,d). We then extended this approach to test the prognostic significance of the target gene signatures of five additional top UR activated in CIMP (CITED2, HIVEP1, IRF2BP2, RNASEH2B and SIRT1) or activated in LOW lesions (TNF, TGM2, STAT1, NFkB and STING1). The target gene signatures of three of these UR (RNASEH2B, TNF, STING1) had prognostic significance in both the EPICA and TCGA MM cohorts (data not shown). Taken together, these findings are consistent with the hypothesis that distinct DNA methylation profiles in MM shape different transcriptional landscapes, which in turn influence clinical behavior.

**Figure 2.**
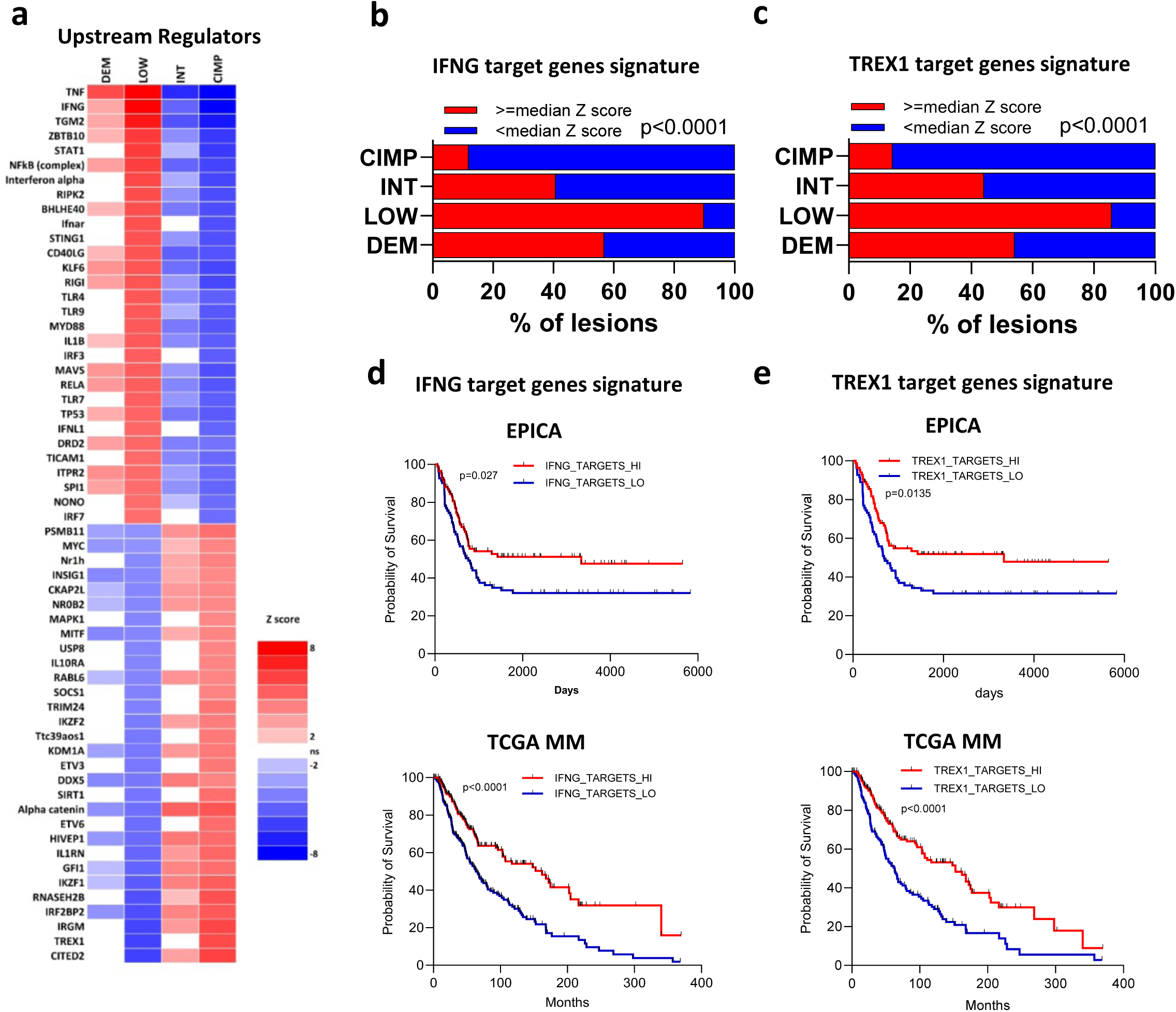
Top master molecules activated in LOW and CIMP MM classes regulate target gene signatures with prognostic significance. **a,** Heatmap of IPA computed Z scores for top UR predicted to be activated (red) or inhibited (blue) in each of the four methylation-defined clusters. **b,c,** Stacked bar plots showing for each methylation cluster in the EPICA cohort the percentage of samples with expression of the IFNG (b) or TREX1 (c) target genes above or below the median z score value of each signature. **d,e,** Kaplan-Meier survival curves of patients in the EPICA cohort (top plot) or TCGA MM cohort (bottom plot) according to median Z score expression (from RNA-seq profiling) of IFNG (d) or TREX1 (e) target gene signatures. In d and e, patients in both cohorts were grouped according to median UR target gene expression above (“HI”) or below (”LO”) the median Z score value of the signature. Statistical analysis b, c by Chi-square; in d, e by log rank test.

### Methylation subsets are associated with ICB response or resistance gene signatures and with clinical outcome after adjuvant immunotherapy

We asked whether the melanoma methylation subsets could be characterized by differential expression of predictive signatures of immunotherapy response. Indeed, two predictive scores, Miracle^27^ and IMPRES^28^ resulted in significantly different values and seven transcriptional signatures^29–33^ were differentially expressed in the four methylation classes of both EPICA and TCGA MM cohorts (Fig. 3a). In the EPICA cohort the viral mimicry signature^34^ and the IFNG ICB response signature^30^, were enriched in the LOW compared to the DEM, INT and CIMP subsets (Fig. 3b-c,e-f). Conversely, the mesenchymal-like (MES) ICB resistance signature^35^ was more expressed in the CIMP and INT subsets compared to the LOW and DEM classes (Fig. 3d,g). The results concerning the latter three signatures were validated in the methylation-defined classes of the TCGA MM cohort (Supplementary Fig. 2a-f). To test the potential relevance of these findings in a therapeutic setting we used RRBS in an independent cohort of n=28 pre-therapy lesions from Stage III-IV melanoma patients treated with adjuvant a-PD-1 and/or a- CTLA-4 ICB (Supplementary Table 4). The EPICA cohort samples were used as training set to classify the pre-adjuvant lesions into the DEM, LOW, INT and CIMP classes (Supplementary Table 4). We performed the subsequent analyses by merging the CIMP and INT, and the LOW and DEM subtypes. Based on the clinical outcome to adjuvant therapy, the CIMP+INT category was enriched among patients experiencing disease relapse (NR in Fig. 3h), although the trend did not reach statistical significance. However, crucially, patients belonging to the DEM+LOW category showed a strong and significant advantage in relapse free-survival compared to those in the INT+CIMP category (Fig. 3i).

**Figure 3.**
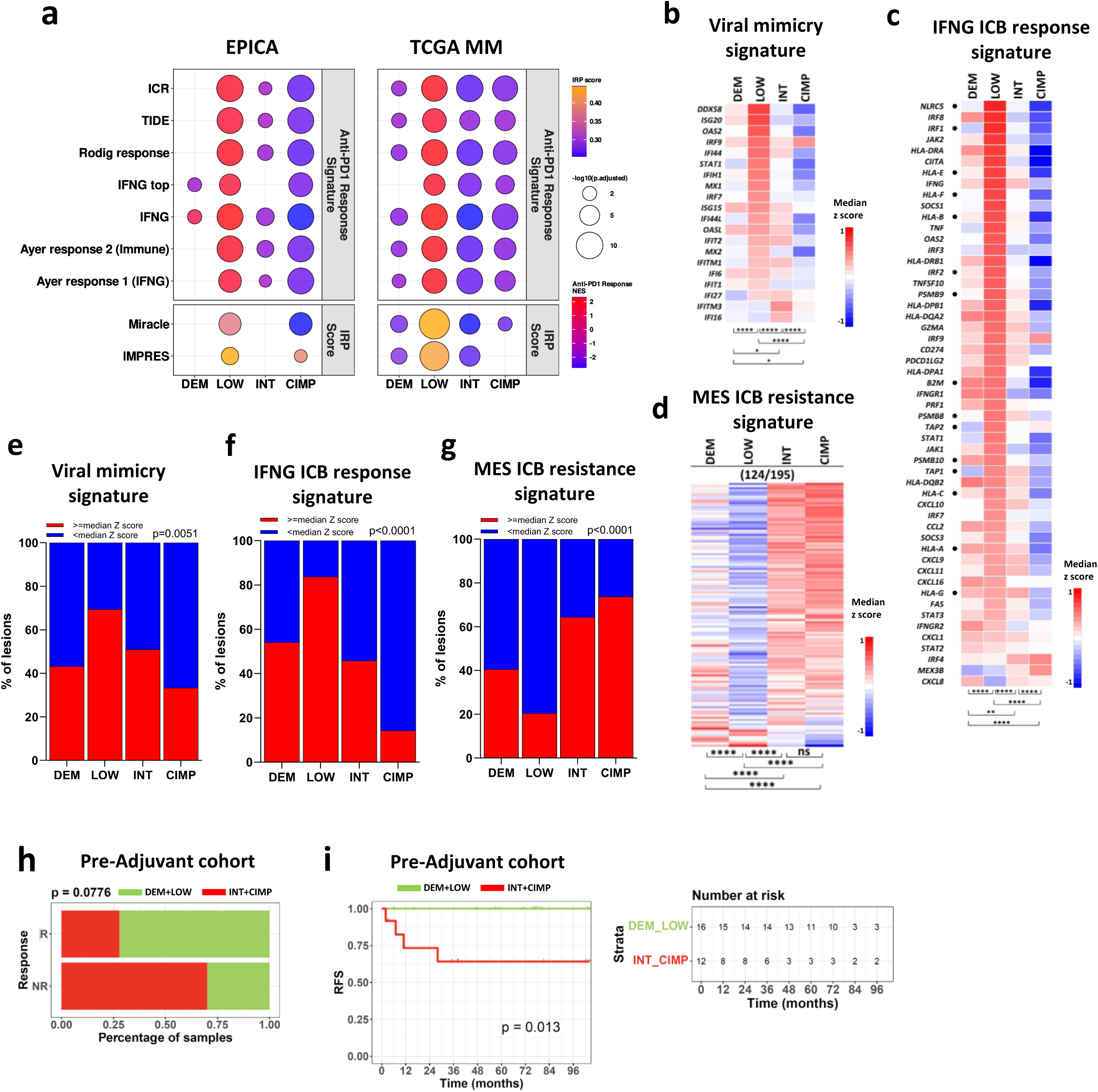
Expression of ICB predictive gene signatures in methylation-defined classes of the EPICA cohort and association of methylation classes with clinical outcome in an adjuvant ICB cohort. **a,** Dot plot of normalized enrichment scores (NES) from GSEA of seven anti-PD1 response signatures in the four methylation subsets of the EPICA and TCGA metastatic melanoma cohorts. Dot size represents the adjusted p-values and scale colors represent the NES (top). Dot plot of immune-response predictor scores (IRP). Dot colors represent the mean score in each group and size the adjusted p-values (bottom). **b-d**, Heatmaps of median Z score expression of genes in the viral mimicry (b), IFN ICB response (c) and MES resistance (d) signatures in the four methylation subsets. Genes identified by dots in panel C represent a core HLA Class I APM gene set. **e**-**g**, Stacked bar plots showing for each methylation cluster in the EPICA cohort the percentage of samples with expression of the viral mimicry (e), IFNG ICB response (f) and MES (g) signatures above or below the median z score value of each signature. Statistical analysis in b-d by Kruskal Wallis test followed by Dunn’s multiple comparison test; in e-g by chi square.*: p<0.05; **: p<0.01, ***: p<0.001; ****: p<0.0001. **h**, Bar plot for patients in an adjuvant ICB therapy cohort showing the percentage of pre-adjuvant samples in DEM/LOW vs INT/CIMP methylation groups, stratified by clinical outcome. Statistical significance by the chi-square test. **i,** Kaplan-Meier relapse-free survival curves for patients in the pre-adjuvant cohort (n=28), stratified by methylation class and grouped into DEM/LOW and INT/CIMP categories. The corresponding risk table is shown on the right.

### An inflamed, T cell rich tumor microenvironment with high frequency of intra-tumor CD8^+^ TCF-1^+^ PD-1^+^ TIM-3^-^ T_PEX_ characterizes LOW lesions compared to CIMP tumors

We used deconvolution algorithms, immunohistochemistry (IHC) and multiplex immunofluorescence to investigate in detail the tumor immune microenvironment of the methylation subsets in the EPICA cohort. Melanoma microenvironment-specific signatures^36,37^ and Xcell algorithm^38^ indicated that the LOW and the CIMP subsets were, respectively, the most and the least enriched for B cells and T cells at distinct stages of functional differentiation, as well as for myeloid or plasmacytoid dendritic cells (Extended Data Fig. 6a,b). Compared to DEM, INT and CIMP subsets, the LOW lesions were enriched for expression of gene signatures defining pro-tumoral macrophages^39^, tertiary lymphoid structures^40^, pre-exhausted T cells (T_PEX_)^41–43^ and exhausted T cells (T_EX_)^43–44^ (Extended Data Fig. 6c-h). These findings were validated in the four methylation classes of the TCGA MM dataset (Extended Data Fig. 6a and data not shown).

We then used IHC to stain EPICA lesions (n=191) for CD3, CD4, CD8, PD-1, PD-L1, FOXP3, CD68 and CD163. The LOW group, when compared to the CIMP subset, showed the highest scores for CD3, CD4, CD8, FOXP3 and PD-1 in intra-tumor and extra-tumor compartments (Fig. 4a). PD-L1 and CD68 had higher IHC scores in LOW vs CIMP lesions in the intra-tumor compartment, (Fig. 4a). The LOW subset was enriched for tumors with a strong intra-tumor infiltrate of CD3^+^ T cells while the CIMP subsets was enriched for lesions with an excluded CD3^+^ and CD8^+^ infiltrate (Extended data Fig. 7a,b). Compared to patients with CIMP tumors, a larger fraction of patients with LOW tumors and with higher-than-median CD3, CD4 and CD8 IHC scores did not undergo further clinical stage progression during follow-up (Supplementary Fig. 3, dark blue bars).

**Figure 4.**
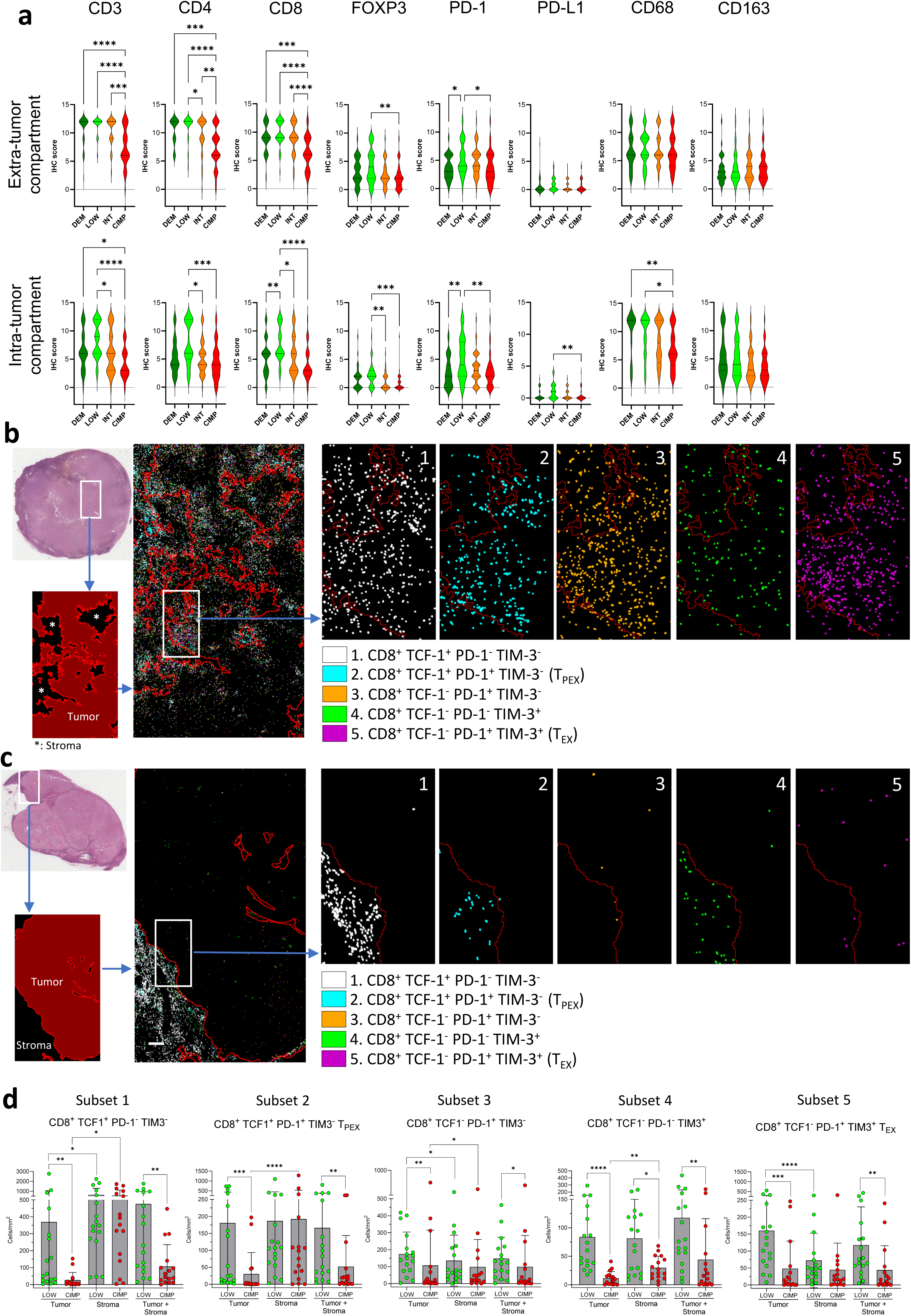
Infiltrating T cells, T_PEX_ and T_EX_ are enriched in LOW melanomas and depleted in CIMP melanomas. **a,** Violin plots showing expression, by semi-quantitative IHC, of CD3, CD4, CD8, PD-1, PD-L1, CD68 and CD163 in extra-tumor or intra-tumor compartments of n=191 EPICA lesions classified according to DEM (n=39), LOW (n=50), INT (n=60), CIMP (n=42) methylation classes. Data expressed as IHC score (see Supplementary Methods). **b**,**c**, Multiplex immunofluorescence analysis of a representative LOW (b) and CIMP (c) lesions. In b and c, the H&E image (top) shows the area used for tissue segmentation (bottom) with tumor and stroma identified in dark red and black, respectively. A higher magnification field of the same area shows the density and position of 5 main CD8^+^ T cell phenotypes identified based on differential expression of TCF-1, PD-1 and TIM-3 and color-coded as indicated. Visualization of tumor cells was omitted in b, c. **d,** Density (cells/mm^2^) in tumor, stroma and whole tissue (tumor + stroma) of the 5 CD8^+^ subsets defined by differential expression of TCF1, PD-1 and TIM-3 in LOW (n=17) and CIMP (n=16) lesions. Statistical analysis: in a by Kruskal Wallis test followed by Dunn’s multiple comparison test; in d, by Mann Whitney test for LOW vs CIMP comparisons in each microenvironment compartment; by Friedman multiple comparison test for tumor vs stroma vs tumor + stroma comparisons within each methylation subset. *: p<0.05; **: p<0.01; ***: p<0.001; ****: p<0.0001.

To identify CD8^+^ subsets at different stages of functional differentiation towards exhaustion in tumor and stroma of EPICA lesions, we looked at expression of CD8, TCF1, PD-1 and TIM-3 by multiplex immunofluorescence analysis in two sets of LOW and CIMP tumors (Supplementary Fig. 4a-d for analysis strategy). This approach allowed to identify 5 CD8^+^ stages, from the less differentiated CD8^+^ TCF1^+^ cells, including the TCF1^+^ PD-1^+^, TIM-3^-^, a profile consistent with T_PEX_^45^, to the more differentiated TCF1^-^, PD-1^+^, TIM-3^+^ phenotype, consistent with T_EX_^46^. All five CD8^+^ subsets including T_PEX_ and T_EX_ (subsets 2 and 5, in Fig. 4b-d) showed significantly higher cell densities in tumor areas of the LOW lesions compared to CIMP lesions (Fig. 4b-d). Taken together, these results indicate that LOW and CIMP lesions have divergent immune contextures, with the LOW subset enriched for T cell inflamed lesions and for presence of CD8^+^ subsets at different stages of functional differentiation, including T_PEX,_ a subset associated with responsiveness to ICB^47^.

### A main mechanism of immune escape is associated with the MM methylation classes

Fifteen genes (black dots close to the gene symbols in Fig. 3c) in the IFNG ICB response signature, represent the core of the HLA class I antigen processing and presentation pathway. A low expression of this gene signature in a tumor lesion may potentially reveal a mechanism of immune escape, i.e. downmodulation/loss of HLA class I molecules on tumor cells. To test this hypothesis, we first set out to identify the most informative genes in the HLA class I pathway. To this end we combined the 15 genes of the IFNG ICB signature with 25 additional genes, selected through literature search, and involved in the HLA class I pathway and in its positive and negative regulation. By spearman correlation analysis of the expression levels we found that 19 out of 40 genes (gene set 2 in Supplementary Fig. 5a, thereafter “HLA-I APM signature”) were directly and significantly correlated in both EPICA and TCGA MM cohorts (Supplementary Fig. 5a). The HLA Class-I APM signature was expressed above median z score values in a higher fraction of LOW compared to CIMP tumors in the EPICA cohort (Supplementary Fig. 5b) and showed prognostic significance in the EPICA and TCGA MM cohorts ((Supplementary Fig. 5c).

We then used IHC, coupled to quantitative digital pathology analysis in the EPICA cohort (n=175 lesions, from n=155 patients) to directly assess expression of HLA Class I molecules on tumor cells. Expression of HLA Class I antigens on tumor cells by IHC and of the HLA Class-I APM gene signature were positively correlated in the whole EPICA cohort (r=0.431, p<0.0001, Fig. 5a). Lesions retaining high expression of HLA Class I or showing variable levels of HLA Class I downmodulation were found in all four methylation subsets (Fig. 5a and Supplementary Fig. 6 for representative stainings). However, lesions expressing HLA class I molecules on >90% of the tumor cells were 51% in the LOW subset, compared to 17.1% in the CIMP lesions (Fig. 5b). Conversely, lesions with almost complete HLA Class I downmodulation (<10% of positive tumor cells) were 29.3% in the CIMP subset compared to 12.2% of the LOW subset (Fig. 5b). In the EPICA cohort, high expression of HLA class I molecules on tumor cells was associated with lack of further AJCC stage progression (Fig. 5c) and with improved survival (Fig. 5d). Moreover, the LOW subset contained the highest proportion of “HLA Class I^HI^ / immune infiltrate^HI^” lesions compared to the CIMP subset (Fig. 5e). Conversely, CIMP subset contained the highest proportion of “HLA Class I^LO^/ immune infiltrate^LO^” lesions compared to the LOW ones (Fig. 5e). Collectively, these data indicate preferential downmodulation/loss of expression of HLA Class I molecules on tumor cells in CIMP compared to LOW lesions and highlight the clinical relevance of HLA Class I downmodulation in MM.

**Figure 5.**
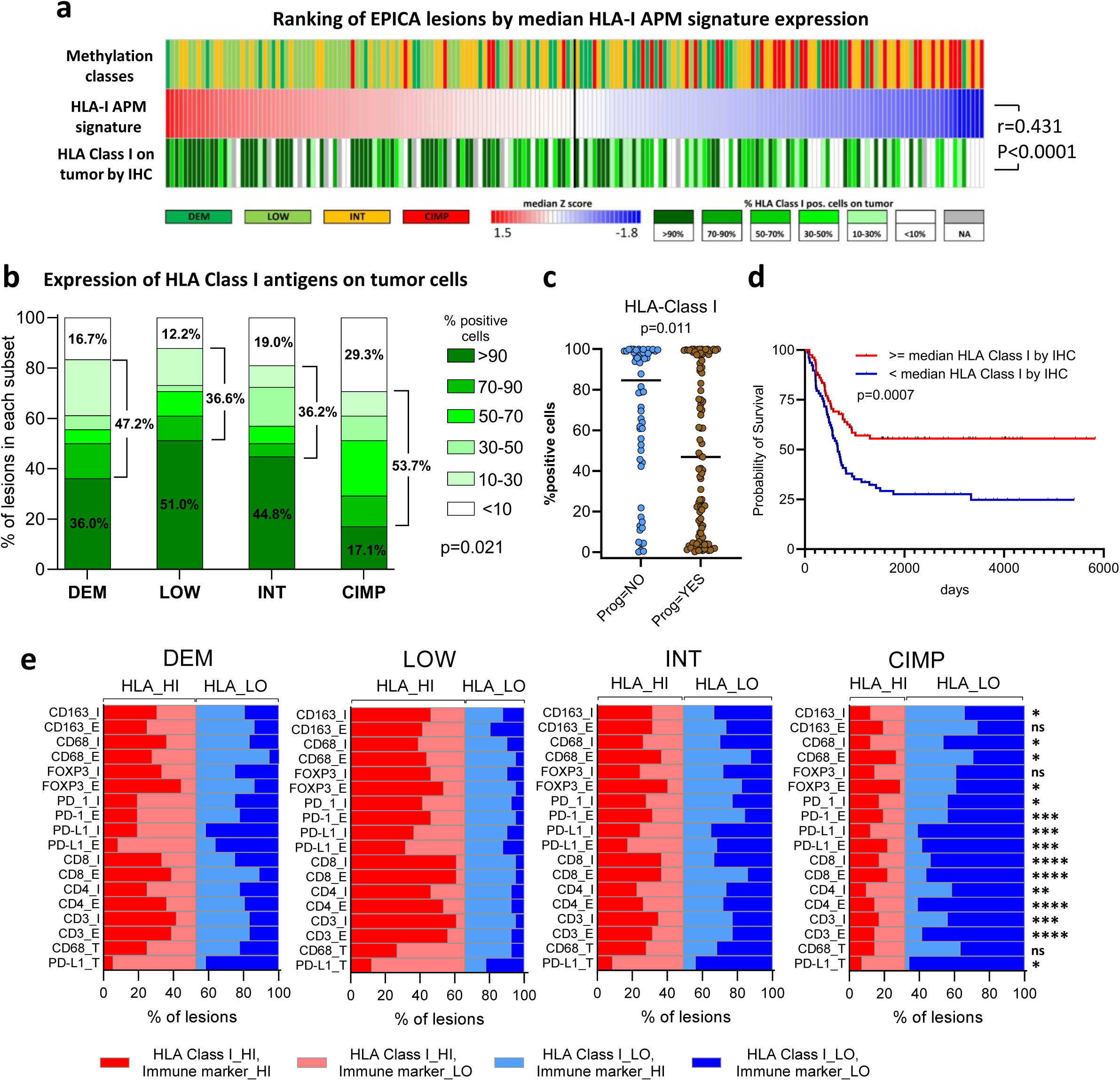
Downmodulation/loss of expression of HLA class I molecules on tumor cells is more frequent in CIMP compared to LOW lesions. **a**, Ranking of EPICA lesions, colored by methylation subset, according to median expression Z score (from RNA-seq profiling) of the HLA Class I APM signature and according to expression of HLA Class I molecules on tumor cells by IHC. **b,** Stacked bar plots showing for each methylation cluster in the EPICA cohort the percentage of samples in each of six classes of HLA Class I antigen expression on tumor cells (by IHC and quantitative digital pathology analysis). **c**, Association of HLA class I antigen expression on tumor cells in the EPICA cohort with progression to any subsequent AJCC stages. **d,** Kaplan Meier survival analysis of patients in the EPICA cohort according to expression of HLA Class I antigens on tumor cells. **e,** Association of methylation classes with immune contexture and with expression of HLA Class I molecules on tumor cells. Expression of HLA Class I and of immune markers, by IHC, was dichotomized into “HI” and “LO” groups corresponding to expression above or below the median value of each parameter. Statistical analysis in a by spearman correlation; in b,e by Chi-square; in c by Mann-Whitney test, in d by log rank test. *: p<0.05; **: p<0.01; ***: p<0.001; ****: p<0.0001.

### CIMP lesions are enriched for proliferative, differentiated melanomas, while LOW lesions are enriched for de-differentiated melanomas

To test the association of methylation subsets with melanoma biological features and differentiation programs we compared expression of four distinct gene signatures in DEM, LOW, INT and CIMP subsets of EPICA cohort. The melanoma-specific cell cycle (MCC) signature^36^ was enriched in the CIMP lesions compared to all other subsets (Extended Data Fig. 8a). The IFNG de-differentiation (IFNG-dediff) signature^48^, unrelated to the IFNG ICB response signature, and the TEADS invasive signature^49^, were strongly enriched in the LOW and DEM subsets compared to the INT and CIMP subsets in both the EPICA and TCGA MM cohorts (Extended Data Fig. 8b-e). We then tested expression in the EPICA methylation subsets of the 7 melanoma differentiation sub-signatures described by Tsoi et al.^50^. The “melanocytic” and “transitory-melanocytic” sub-signatures were more expressed in CIMP and INT tumors. The “neural crest-like”, “undifferentiated/neural crest-like” and “undifferentiated” sub-signatures were more expressed in the LOW compared to the CIMP lesions (Extended Data Fig. 8f). Taken together, these data indicate that the DNA methylation class is associated with the melanoma differentiation phenotype: CIMP tumors are enriched for differentiated, proliferative melanomas, while LOW lesions are enriched for de-differentiated melanomas.

### The differentiation and immune-related transcriptional profiles observed in CIMP vs LOW lesions are tumor-intrinsic programs retained in-vitro by melanoma cell lines

We used a panel of 46 melanoma cell lines (n=45 from metastatic lesions, n=1 from a primary tumor) derived from surgical specimens of patients (Supplementary Table 5 for cell line origin and STR profiling authentication). By qPCR for two markers of differentiated melanomas (MITF and PMEL) we identified MITF/PMEL^HI,^ MITF/PMEL^INT^ and MITF/PMEL^LO^ cell lines (Fig. 6a). Transcriptomic analysis revealed higher expression of the “melanocytic” and “transitory-melanocytic” sub-signatures^50^, in the MITF/PMEL^HI^ cell lines (PLN74 and GML41), and higher expression of the “neural crest-like”, “undifferentiated-neural crest-like” and “undifferentiated” sub-signatures in two MITF/PMEL^LO^ cell lines (BRM17 and VRG100, Fig. 6b). Consistently, expression of the TEADS invasive signature was highest in the MITF/PMEL^LO^ cell lines compared to MITF/PMEL^MID^ and MITF/PMEL^HI^ cell lines (Fig. 6c). Crucially, the viral mimicry and the IFNG-ICB response signatures were expressed at higher levels in the de-differentiated MITF/PMEL^LO^ cell lines compared to the differentiated MITF/PMEL^HI^ cell lines (Fig 6d,e), mirroring the different profiles observed in LOW vs CIMP lesions, respectively.

**Figure 6.**
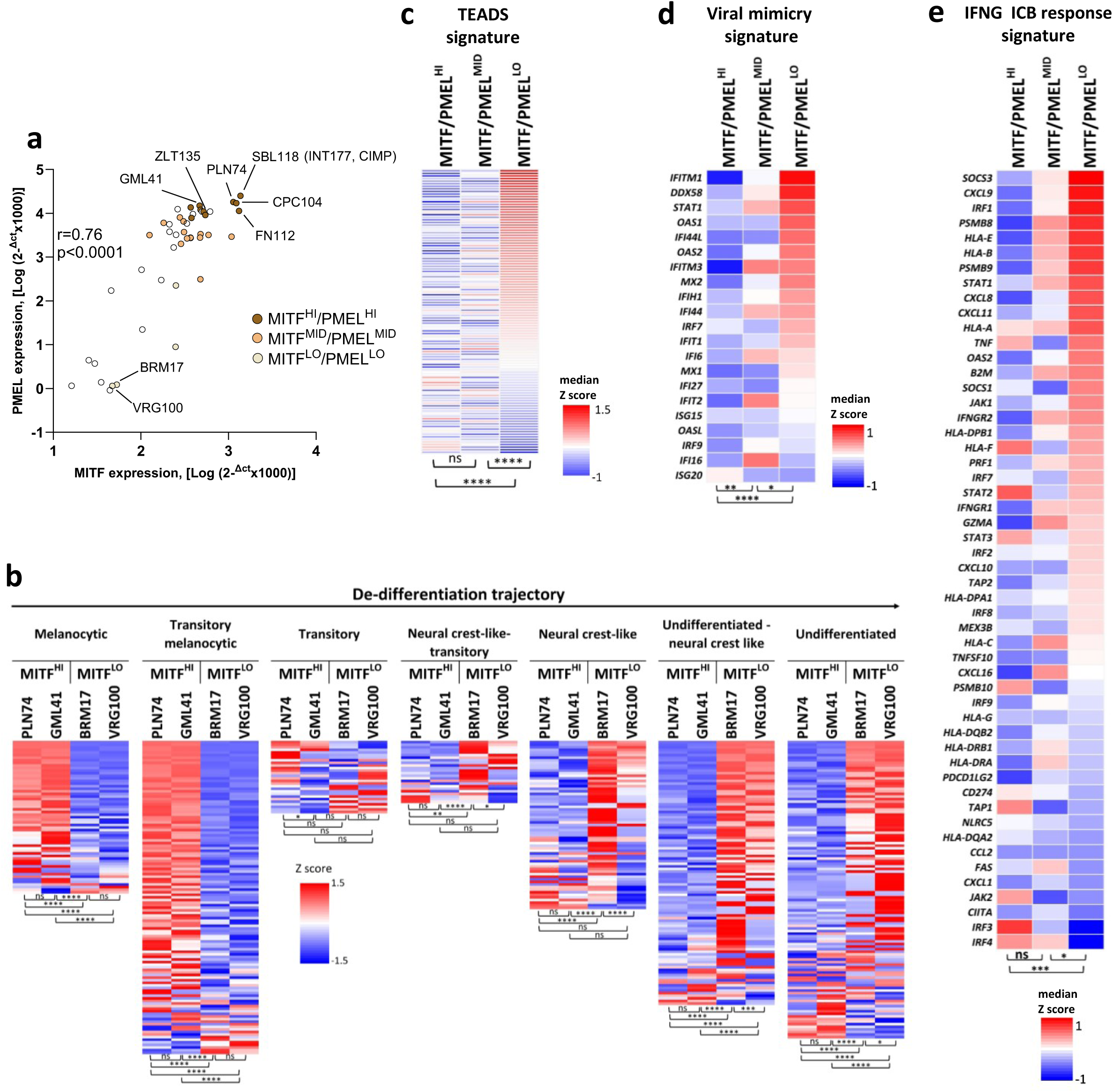
Differential expression of melanoma differentiation signatures and of immunotherapy response signatures in MITF/PMEL^HI^ vs MITF/PMEL^LO^ cell lines. **a**, Expression of MITF and PMEL genes, by qPCR, in 46 melanoma cell lines. Subsets of cell lines with high (HI), intermediate (MID) and low (LO) MITF/PMEL expression are highlighted by the indicated color code. Cell line SBL118 was generated from the surgical specimen of the lesion corresponding to sample INT177 of the EPICA cohort. **b**, Heatmaps showing expression, from Clariom S array data, in four melanoma cell lines of the 7 melanoma differentiation sub-signatures described by Tsoi et al.^50^. **c**,**d**,**e**, Heatmaps showing expression, from Clariom S array data, in the three sets of melanoma cell lines (MITF/PMEL^HI^, MITF/PMEL^MID^, MITF/PMEL^LO^) of the TEADS invasive signature (c), of the viral mimicry (d) and of the IFNG-ICB response signatures (e). Statistical analysis in a by spearman correlation; in b by two-way anova followed by Tukey’s multiple comparison test; in c, d, e by Kruskal Wallis test followed by Dunn’s multiple comparison test. *: p<0.05; **: p<0.01; ***: p<0.001, ****: p<0.0001.

### Treatment of differentiated melanoma cell lines with a DNMTi induces global DNA de- methylation, promotes de-differentiation and upregulates ICB predictive signatures

Six MITF/PMEL^HI^ differentiated melanoma cell lines were treated with the DNMTi guadecitabine for 7 to 21 days, according to a previously published protocol^20^. Based on the extent of global DNA de- methylation, measured by EPIC arrays in guadecitabine-treated vs controls, four out of six cell lines (CPC104, ZLT135, GML41, PLN74) were highly responsive to this DNMTi, (Fig. 7a). In the responsive cell line PLN74, global DNA de-methylation promoted by guadecitabine increased at 14 and 21 days, compared to 7 days of treatment (Fig. 7a). Two cell lines (SBL118 and FN112) were weakly responsive to the DNMTi, as they showed only modest reduction in global DNA methylation levels at day 7 of treatment, with no further decrease at 14 and 21 days (cell line FN112, Fig. 7a). By analysis of Clariom S transcriptomic data in the highly responsive PLN74 cell line, we found that genes in the proliferative MCC signature^36^ and in “melanocytic” and “transitory-melanocytic” sub-signatures^50^ were downmodulated by guadecitabine treatment (Extended Data Fig. 9a). In contrast, genes in the TEADS^49^ and IFNG-dediff signatures^48^, as well as genes in the “neural crest-like”, “undifferentiated-neural crest-like” and “undifferentiated” sub-signatures^50^ were progressively upregulated (Extended Data Fig. 9a). In the same PLN74 cell line, guadecitabine treatment strongly downmodulated expression of three differentiation proteins (MITF, GP100/PMEL, MART-1), as evaluated by quantitative digital pathology on melanoma cell cytospins (Extended Data Fig. 9b,d). In contrast, markedly reduced effects were observed on the expression of these three markers in the poorly responsive FN112 cell line (Extended Data Fig. 9c,e). We then tested guadecitabine effects on melanoma cell clones isolated from the same metastatic tumor 665/2 by cloning in soft agar and micromanipulation^51^. Based on the Tsoi et al. differentiation sub-signatures^50^ clones 2_21, 2_33 and 2_59 exhibited a differentiated profile, while clones 2_4, 2_14 and 2_17 showed a predominant de-differentiated profile (Supplementary Fig. 7a). Upon treatment for 7 days with guadecitabine, the differentiated melanoma clone 2_59 underwent a process of de-differentiation (Supplementary Fig. 7b), as shown by downmodulation of “melanocytic” and “transitory-melanocytic” sub-signatures and by upregulation of “neural crest-like”, “undifferentiated-neural crest-like” and “undifferentiated” sub-signatures^50^. No consistent profile shift was observed in the de-differentiated clone 2_17.

**Figure 7.**
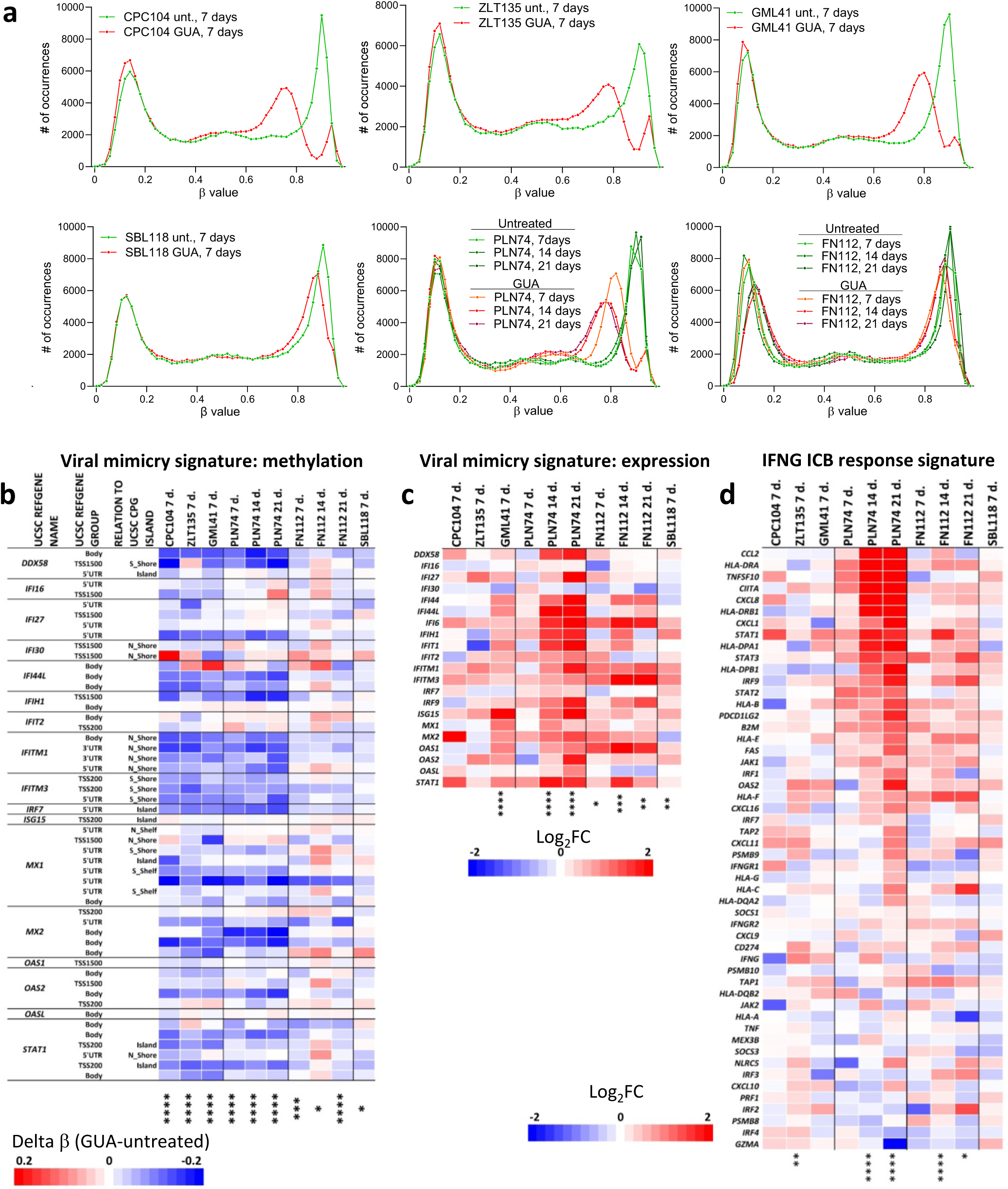
DNA de-methylation, promoted by a DNMTi, shifts the transcriptional profile of differentiated melanoma cell lines towards the “immune-high” phenotype found in LOW lesions. **a**, Global DNA methylation profiles, from EPIC array data, of six differentiated melanoma cell lines untreated or treated with guadecitabine (GUA) for 7 to 21 days. **b**, Changes in methylation level induced by guadecitabine treatment at methylation sites corresponding to genes in the viral mimicry signature in 6 differentiated melanoma cell lines cultured with guadecitabine for 7 to 21 days. **c**,**d**, Treatment with guadecitabine for 7 to 21 days of six melanoma cell lines modulates genes in the viral mimicry signature (c) and in the IFNG ICB response signature (d). Statistical analysis in c-e by Mann-Whitney test. *: p<0.05; **: p<0.01; ***: p<0.001; ****: p<0.0001.

We then evaluated the impact of guadecitabine treatment on the methylation and expression of genes belonging to immunotherapy response signatures. Strong de-methylation at sites corresponding to genes in the viral mimicry signature was observed (Fig. 7b) in four highly responsive cell lines (CPC104, ZLT135, GML41, PLN74). Upregulation of genes in this signature was also observed in four cell lines and most significant effects were induced in PLN74 cells at day 14 and day 21 of treatment (Fig. 7c). In 3/6 melanoma cell lines significant upregulation of the genes in the IFNG-ICB response signature was also observed upon guadecitabine treatment (Fig.7d).

Finally, we selected the top UR found constitutively activated or inhibited in CIMP lesions (shown in Fig. 2a) and tested whether the functional status of these UR could be reversed in differentiated melanoma cell lines (PLN74, GML41, CPC104, FN112 and SBL118) upon treatment with guadecitabine. In all these cell lines we found that DNMTi treatment activated the UR found inhibited in CIMP tumors and inhibited the UR found activated in CIMP lesions (Extended Data Fig.10). Taken together, these results indicate that global DNA de-methylation, induced by a DNMTi, profoundly reshapes the transcriptomic profile of differentiated melanoma cell lines towards a de-differentiated and “immune-high” phenotype mirroring that observed in LOW tumors.

## Discussion

In this study, by exploiting epigenetic variability at the methylome level, we identified four methylation subsets in the EPICA and TCGA MM cohorts. Extensive multi-omics characterization of the four subsets revealed that the epigenetic classification provides a framework for explaining several layers of MM complexity including: transcriptional programs of melanoma differentiation and immunity, prognostically relevant gene signatures controlled by master molecules, expression of ICB response and resistance signatures, specific immune contexture structure, a main immune escape mechanism, propensity for subsequent stage progression and clinical outcome. The LOW and the CIMP tumors exhibited the most divergent phenotypic profiles and appear to represent opposite ends of the spectrum of melanoma biological heterogeneity. The enrichment, in tumors of the LOW class, for prognostically relevant signatures and for lesions with a strong T cell infiltrate, suggests that this subset contains a high proportion of metastatic lesions with an active anti-tumor response. Such immune response may contribute to reduce the likelihood of further clinical stage progression and to improve overall survival, as we found in patients with LOW tumors compared to patients with CIMP lesions. The high expression of ICB predictive signatures and the enhanced density at tumor site of CD8^+^ T cell subsets known to be associated with response to immunotherapy (such as the T_PEX_), suggests that patients with LOW lesions might have increased responsiveness to ICB. In support of this hypothesis, in an independent adjuvant ICB cohort, we found that patients with DEM/LOW pre-therapy lesions had a significantly better relapse-free survival compared to patients with INT/CIMP lesions. These findings suggest that the DNA methylation profile of neoplastic lesions could be exploited to predict which MM patients may benefit or show resistance to ICB therapy. Indeed, at least two recent studies^52,53^ in the metastatic setting support this conclusion. In one of these studies^53^, three methylation subsets were identified. Cluster 1, the most hypomethylated, contained only responders, cluster 2 contained 50% of responders while the hypermethylated cluster 3 contained 39 non-responders out of 45 patients in the cluster.

Compared to LOW lesions, CIMP tumors showed a differentiated, proliferative profile, higher expression of an ICB resistance signature, higher frequency of lesions with HLA Class I downmodulation/loss on neoplastic cells, a frequently cold immune contexture and worse overall survival. In agreement with our findings, one of three methylation subtypes identified among 50 MM by Lauss et al.^15^ showed elevated methylation at promoter islands and consisted of “proliferative” and “pigmentation type” tumors, thus resembling the differentiated CIMP subset defined in this study. The evidence for worse clinical outcome in the CIMP vs the LOW class, that we found in both the EPICA and TCGA MM cohorts, is to be considered in the light of similar evidence obtained by Conway et al in primary melanomas^14^: collectively these findings suggest that acquisition of the hypermethylated profile “imprints” a poor clinical outcome across all stages of melanoma progression. In agreement, in the recent study^17^ the authors found a worse outcome in Stage II and III melanoma patients whose primary tumors were classified in the CIMP methylation class.

The experiments with a panel of melanoma cell lines allowed us to test the cause-effect relationship linking the global DNA methylation profile of melanoma cells to their transcriptional programs. First, we found that differentiated and de-differentiated melanoma cell lines showed respectively lower vs higher expression of the viral mimicry and ICB response signatures. These findings suggested that level of constitutive expression of clinically relevant immune-related signatures is a cell autonomous feature of melanoma cells associated with their melanocyte differentiation programs. Second, promotion of global DNA de-methylation by the DNMTi guadecitabine shifted the phenotype of differentiated melanoma cells towards a de-differentiated and “immune-high” phenotype. Taken together these findings support the notion that global DNA methylation profile of MM cells contributes to shape the overall transcriptional phenotype of the tumor. Moreover, these findings provide a pre-clinical rationale supporting use of demethylating agents to promote *in-vivo* the rewiring of melanoma transcriptional programs towards an immunotherapy responsive profile. Results of the multi-omics analysis of pre- and post-therapy lesions from MM patients enrolled in our Phase 1b NIBIT-M4 trial with guadecitabine + ipilimumab support this notion^54^. Although that trial was not designed to compare the effect of a demethylating agent plus ICB vs ICB alone, post-therapy lesions from responding patients showed promotion of expression of ICB response signatures and a significant anti-correlation between methylation and expression of LINE, SINE and LTR elements, including ERVs^54^. These data suggest that a demethylating agent may, through global demethylation, reshape the transcriptional programs of melanoma cells in-vivo and activate transcription of ERV sequences, that in turn trigger the viral mimicry process crucial for ICB response^34^.

## Materials and Methods

### Patients

Tumor samples (n=191) from the EPICA cohort of MM lesions were obtained based on informed consent from n=165 Stage III and IV patients according to AJCC 8^th^ edition. The study was conducted according to the Declaration of Helsinki Principles and following approval by the Ethics Committee of Fondazione IRCCS, Istituto Nazionale dei Tumori, Milan, Italy (protocol number INT 170/18). Patient and lesion eligibility criteria are described in Supplementary Methods. Relevant demographic and clinicopathological data of the EPICA cohort are listed in patient centric form in Supplementary Table 1a and in lesion centric form in Supplementary Table 1b. Tumor samples (n=28) from an additional cohort of MM patients (Supplementary Table 4) were obtained from n=28 Stage III or IV-resected (according to AJCC 8^th^ edition) patients treated with adjuvant ICB therapy in daily practice or within clinical trials (CA-209-238 (NCT02388906)^55^; CA-209-915 (NCT03068455)^56^ at the Center for Immuno-Oncology, Department of Oncology, University Hospital of Siena, Siena, Italy. All patients provided an informed consent.

### DNA extraction from FFPE sections and quality controls

Isolation of total DNA from FFPE sections was performed by using Maxwell (R) RSC FFPE Plus DNA Kit (Promega, Madison, WI, USA) according to the manufacturer’s protocol. Quality controls (QCs) were performed on the TapeStation 4200 system using the Genomic DNA High Sensitivity ScreenTape Assay (Agilent Technologies, Santa Clara, CA, USA). All samples had a concentration range between 5-10 ng/µl and a DNA integrity number (DIN) between 2 and 5. Library preparation protocols were adapted to optimize the result obtained with low integrity samples. Isolated DNA (300 ng) was exploited to perform further methylation analyses. Briefly, Ovation® RRBS Methyl-Seq (Tecan/NuGEN, Redwood City, CA) was used for library preparation following the manufacturer’s instructions. Unmethylated lambda phage DNA was spiked-in to estimate the bisulphite conversion rate. DNA samples were quantified with Qubit 2.0 Fluorometer (Invitrogen, Carlsbad, CA). Final libraries were checked with both Qubit 2.0 Fluorometer (Invitrogen, Carlsbad, CA) and Bioanalyzer HS DNA assay (Agilent technologies, Santa Clara, CA). Libraries were then prepared for sequencing and sequenced on paired-end 150 bp mode on NovaSeq6000 (Illumina, San Diego, CA).

### Whole-Exome Sequencing (WES)

WES was performed using Human Comprehensive Exome kit (Twist Bioscience, San Francisco, CA, USA) starting from 200 ng total input. Samples with DIN < 3 were fragmented for 4 minutes at 32°C to maintain 200 bp. Half-volume hybridization was performed at a temperature of 62°C in multiplexes of 12 samples to increase complexity when quality control of the indexed libraries was < 100 ng total per sample. Quality controls and WES data analysis pipeline is described in Supplementary Methods.

### RRBS sequencing

Reduced representation bisulphite sequencing (RRBS) raw reads were trimmed for adaptor sequences using trim galore (v. 0.6.5) (http://www.bioinformatics.babraham.ac.uk/projects/trim_galore/) and filtered for low-quality sequences using fastQC (v.0.11.8) (https://www.bioinformatics.babraham.ac.uk/projects/fastqc/). High quality trimmed reads were mapped to the Human reference genome (UCSC genome assembly GRCh38/hg38) using Bismark (v.0.22.3)^57^ with default parameters. Methylation data as β values for CpG sites, promoters, and genes were retrieved from Bismark coverage outputs using R package RnBeads 2.0 (v.2.6.0) with default parameters^58^. RRBS data analysis process is described in Supplementary Methods.

### Methylation analysis by Infinium MethylationEPIC

Microarray analyses was performed by Genomix4life S.R.L. (Baronissi, Salerno, Italy). DNA concentration in each sample was assayed with a fluorimeter Qubit Fluorometer 4.0 (Invitrogen Co., Carlsbad, CA), Nanodrop One (Thermo Fisher) and its quality assessed with the TapeStation 4200 (Agilent Technologies). Bisulphite converted DNA (250 ng) was used for analysis of whole-genome methylation using the Infinium MethylationEPIC v2.0 kit (Illumina, San Diego, CA, USA), which contains ∼930K unique methylation sites in the most biologically significant regions of the human methylome. In brief, bisulphite converted DNA was whole-genome amplified for 20 h followed by end-point fragmentation. Fragmented DNA was precipitated, denatured and hybridised to the BeadChips for 20 h at 48 °C. The BeadChips were washed and the hybridised primers were extended and labelled before scanning the BeadChips using the Illumina iScan system. GenomeStudio software (version 2011.1; Illumina Inc.) was used for the extraction of DNA methylation signals from scanned arrays. The methylation level for each cytosine was expressed as a beta value calculated as the fluorescence intensity ratio of the methylated to unmethylated versions of the probes: beta values ranged between 0 (unmethylated) and 1 (methylated).

### RNA extraction from FFPE sections and quality controls

Total RNA was isolated from two to four 12 μm FFPE sections by RecoverAll Total Nucleic Acid Isolation Kit (Invitrogen), according to the manufacturer’s protocol including ethanol precipitation. RNA quantity and purity were estimated by a Nanodrop 2000 spectrophotometer (Thermo Fisher Scientific). Further quality controls (QCs) were performed on the TapeStation 4200 system using the RNA High Sensitivity ScreenTape Assay (Agilent Technologies, Santa Clara, CA, USA). All samples had a concentration range between 5-10 ng/µl and RNA integrity number (RIN) between 2-5. The RNA exome analysis using sequence-specific capture of the coding regions of the transcriptome were accomplished by Illumina RNA Prep with Enrichment (Illumina, San Diego, CA, USA). To provide high reproducible sample handling RNA libraries were generated with epMotion^®^ 5075 Liquid Handler (Eppendorf, Hamburg, Germany). All samples were sequenced on an Illumina NovaSeq 6000 sequencer in paired-end mode, generating 100 nt length reads, to obtain an average of 60 million clusters for RNA. Demultiplexing was performed using Illumina bcl2fastq2.

### RNA sequencing

Fastq quality was assessed using fastQC (v. 0.11.8) (https://www.bioinformatics.babraham.ac.uk/projects/fastqc/) and low-quality reads were discarded. Sequence reads were aligned to Human reference genome (UCSC genome assembly GRCh38/hg38) using STAR (v. 2.7.0b)^59^, and the expression was quantified at gene level using featureCounts (v. 1.6.3), a count-based estimation algorithm^60^. Downstream analysis was performed in the R statistical environment as described in Supplementary Methods.

### Quantitative PCR analysis

Total RNA (0.5μg) was reverse transcribed with oligo d(t) using Superscript IV Reverse Transcriptase II (Invitrogen). Real-time PCR, in duplexing method, was carried out with 20ng input cDNA, 1X TaqMan Gene Expression Assays and Taqman Fast Advanced Master Mix on a Quant Studio 1 Real-Time PCR System (Applied Biosystems). The data were analysed by automated baseline and threshold determination. Relative expression was determined on triplicate reactions using the formula 2^-ΔCt^, reflecting target gene expression normalized to endogenous control level (GAPDH and GUSB). The following TaqMan assays were used: GAPDH_fam: GGGCGCCTGGTCACCAGGGCTGCTT (Hs99999905_m1); GUSB_vic: TGAACAGTCACCGACGAGAGTGCTG (Hs99999908_m1); MITF_fam: TCACAGAGTCTGAAGCAAGAGCACT (Hs01117294_m1); PMEL_vic: CATCTCTGATATATAGGCGCAGACT (Hs00173854_m1).

### Melanoma cell lines and treatment with guadecitabine

Melanoma cell lines (n*=*46) were established, maintained and routinely tested for the absence of mycoplasma contamination by PCR, as previously described^61^. The tissue of origin of the cell lines and cell line authentication by STR profiling (Gene-Print10 kit, Promega) are described in Supplementary Table 5. Melanoma cell lines were seeded at 1.25 × 10^4^/mL in T75 flasks (Greiner Bio-One) with RPMI-1640 medium (Life Technologies Limited) supplemented with 4% FCS (Biological Industries) without antibiotics, and were treated with DNMTi guadecitabine (MedChemExpress) according to a previously published protocol^20^. Briefly, cells were seeded at day 1. Guadecitabine was added at 1000 nM at days 2, 4, 8, 11, 15 and 18. Cells were harvested for gene expression and methylation profiling at day 7, or at day 14, or at day 21.

### Immunohistochemistry

Immunohistochemistry on formalin-fixed, paraffin-embedded (FFPE) tissues from human melanoma lesions was performed as described previously^62^. Briefly, 3 μm thick sections were stained with either Leica Bond RX stainer (Leica Biosystems, Buffalo, IL, USA) or Ventana BenchMark ULTRA immunostainer (Ventana Medical Systems, Tucson, AZ, USA). The following antibodies were used: CD3 (Ventana Medical System, clone 2GV6), CD4 (Ventana Medical System, clone SP35). CD8 (Ventana Medical System, clone SP57), CD68 (Dako Agilent, clone KP1), CD163 (Novus Biologicals, clone MRQ-26), Fox-P3 (Abcam, clone 236A/E7), PD-L1 (Dako Agilent, clone 22C3), PD-1 (Biocare Medical, clone NAT105), Beta-2-Microglobulin (Dako Agilent, Polyclonal) and HLA-Class I (Abcam, clone EMR8-5). EMR8-5 recognizes HLA Class I A,B,C and E antigens^63^. IHC data analysis is described in Supplementary Methods.

### Multiplex immunofluorescence (mIF) analysis of tumor microenvironment

Staining of 4-5 μm thick sections from FFPE sections was performed using the Leica Bond RX stainer (Leica Biosystems, Buffalo, IL). The Bond RX staining protocol was implemented as described by Parra et al.^64^. Briefly, after baking and dewaxing with Bond Dewax solution (Leica Biosystems), slides were subjected to heating at 95°C for 20 min in the presence of Bond Antigen Retrieval Tris-EDTA buffer or citrate buffer, depending on the markers. Slides were then incubated with the primary antibodies for 30 min. The following antibodies were used: CD8 (Leica Biosystems, clone 4B11), TCF1/TCF7 (Cell Signalling, clone C63D9), TIM3 (Cell Signalling, clone D5D5R) as well as PD-1 (clone EPR4877(2)) and S100/SOX10 (Clone EP268 1/4C4.9) ready-to-Use antibodies all included in the Opal 6-Plex detection kit (Akoya Biosciences, Marlborough, MA, USA). DAPI was included for identification of nuclei. After wash and incubation with HRP following anti-mouse or anti-rabbit secondary antibodies slides were incubated with fluorophore tyramides of the Opal 6-Plex Detection Kit (Akoya Biosciences) containing the fluorophores Opal 480, Opal 520, Opal 620, Opal 690, Opal 780. After subsequent washes with Bond wash solution slides were counterstained with DAPI, to visualize nuclei. Slides were finally mounted with ProLong Diamond Antifade Mountain (Invitrogen/Thermo Fisher Scientific, Waltham, MA). mIF data analysis is described in Supplementary Methods.

### Statistical analysis

Correlations between methylation subtypes and pathological variables were analysed using Pearson’s Chi-squared Test. The Chi squared test was also used to compare the proportion of cases in each methylation class classified by the level of T cell infiltrate, stage progression expression above or below the median z score value of gene signatures of interest, expression of immune markers and HLA class I antigens on tumor cells. The Kruskal Wallis test followed by Dunn’s multiple comparison test was used to compare methylation classes for expression of median z score values of gene signatures of interest. The Student T test was used to compare the expression of MITF, HMB45 and MART-1 proteins in cell lines treated or not with guadecitabine. Two-way ANOVA and Tukey’s multiple comparison test was used to compare the expression of melanoma differentiation signatures and the effects of guadecitabine treatment in melanoma clones. Survival curves were estimated using the survival R package (v. 3.2–10) and plotted using the Kaplan–Meier method, implemented in the survminer (v. 0.4.9) R package. Log-rank tests were used to compare curves between groups. Results of the multivariable Cox proportional hazards model on the TCGA skin cutaneous melanoma dataset were obtained through the outcome module of the TIMER2.0 web server (available at http://timer.comp-genomics.org)^26^. Differentially expressed genes for UR analysis among methylation-defined subsets were identified by BRB array tools with the following criteria: gene-level p value: 0.001; FDR<0.01; permutation p value (based on 10,000 permutations): <0.01. Lists of differentially expressed genes were used for UR analysis by Ingenuity Pathway Analysis, as described in Supplementary Methods.

## Supporting information

Source data to main figures

## Acknowledgements

The research leading to these results has received funding from Fondazione AIRC under 5 per Mille 2018 - ID.21073 project – P.I. Maio Michele, G.L. Anichini Andrea, G.L. Ceccarelli Michele, G.L. Daniela Massi. This work was also supported by the Ministry of Health, Lombardy and Tuscany Regions, Bando Ricerca Finalizzata, grant number NET-2016-02361632, Program P.I. Michele Maio, WP2 P.I. Andrea Anichini.

## Data availability

Raw data will be deposited on EGA and GEO before acceptance.

## Author contributions

Conceptualization: A.A., A.M.D.G., D.M., M.Milione, B.V., A.M., R.P., M.S., M.Maio, M.Ceccarelli, R.M. Clinical database management: A.A., A.M.D.G., E.S., M.Milione, A.M., R.P., M.S., M.Maio, R.M. Patients’ management: A.M.D.G., A.M., R.P., M.S., M.Maio. Patients’ sample collection: V.L., G.N., A.M., I.B., A.C., M.F.L., S.C., E.S., B.V., M.C., F.U., S.S, M.Milione. Design of experiments: A.A., V.L., A.C., M.F.L., S.C., B.V., M.C., F.U., S.S., D.M., M.Milione, R.M. Performing experiments: G.N., A.M., I.B., V.L., F.S., A.C., M.F.L., F.U., S.S. Bioinformatics analyses: F.P.C., T.M.R.N., R.T., M.Ceccarelli. Data analysis: A.A., F.P.C., V.L., F.S., A.C., M.F.L., S.C., A.M.D.G., E.S., B.V., M.C., F.U., S.S., M.Milione, A.M., M.Ceccarelli, R.M. Technical assistance: G.N., A.M., I.B. Manuscript writing: A.A., F.P.C., T.M.R.N., A.C., M.F.L., A.M.D.G., A.M., R.P., M.S., M.Maio, M.Ceccarelli, R.M. Project supervision: A.A., D.M., M.Maio, M.Ceccarelli, R.M. All authors commented on the manuscript.

---

**Extended Data Figure 1.**
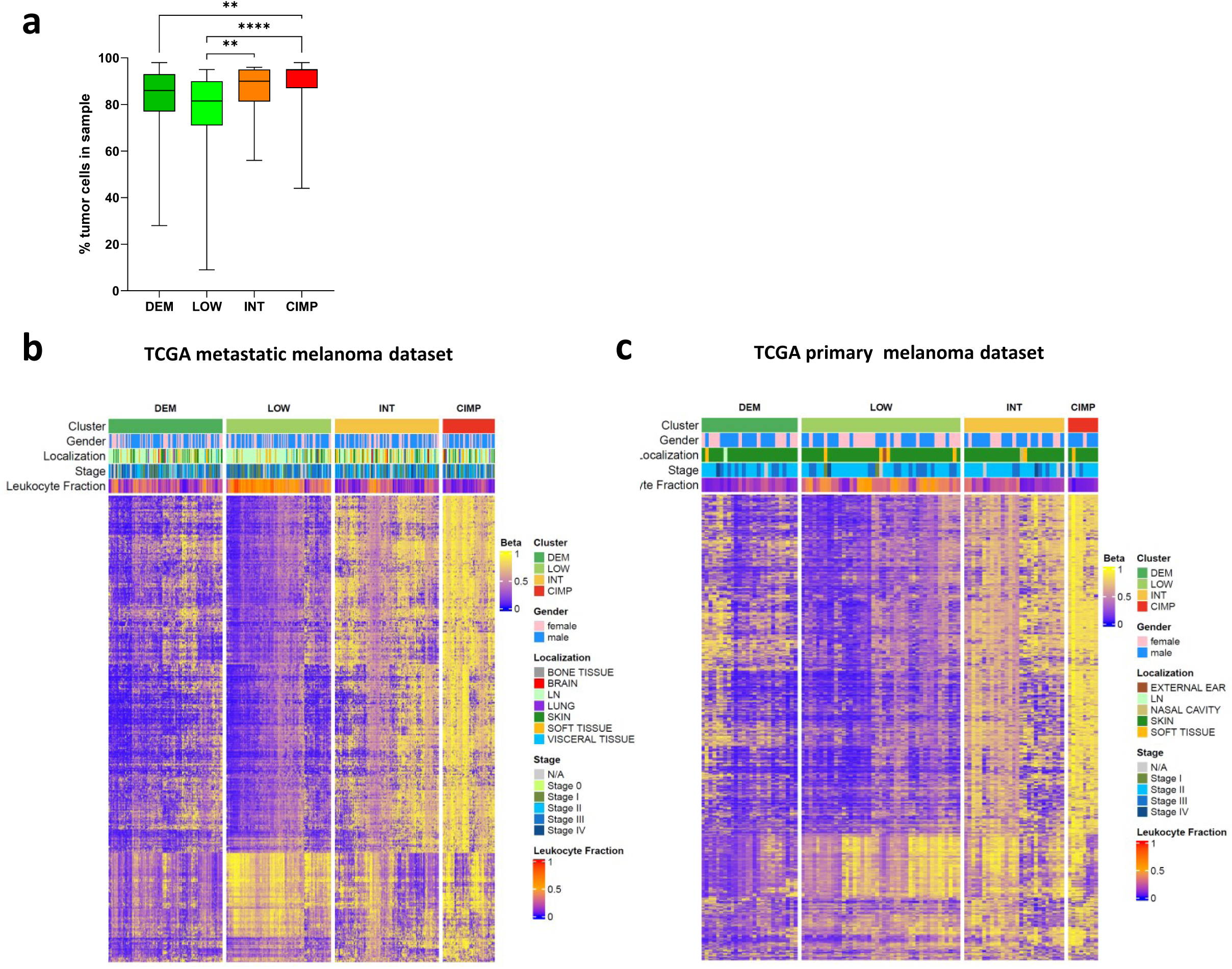
Tumor purity in EPICA methylation subsets and validation of methylation classes in Firehose Legacy TCGA primary and metastatic melanoma cohorts. **a,** Tumor purity in the 191 EPICA lesions classified for the methylation subset. **b,** Unsupervised clustering of n=368 TCGA metastatic SKCM based on 3515 most variable CpG probes. **c,** Unsupervised clustering of n=104 primary melanomas cohort from TCGA based on 5605 most variable CpG probes. Statistical analysis in A by Kruskal Wallis test followed by Dunn’s multiple comparison test.

**Extended Data Figure 2.**
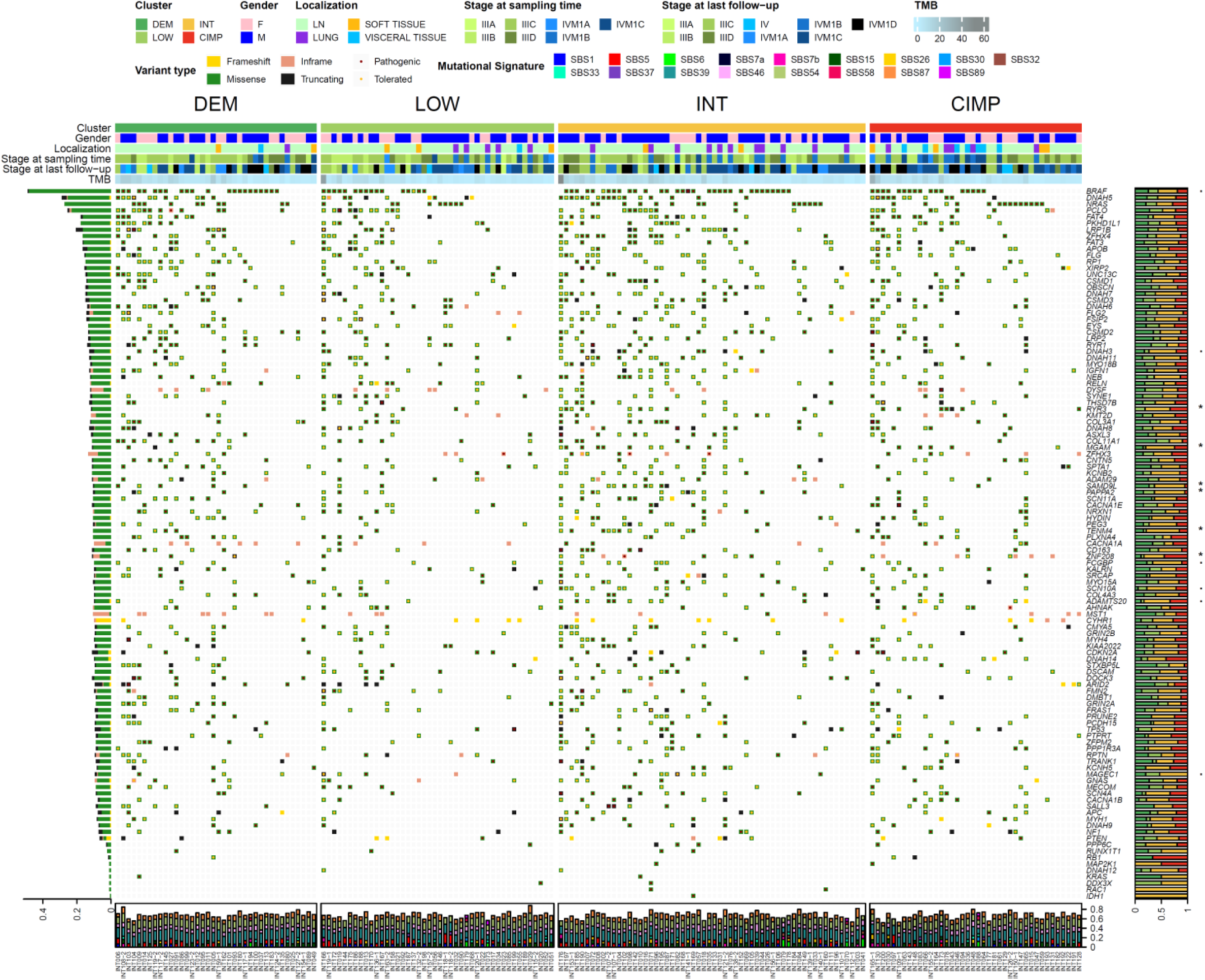
Oncoplot displaying common somatic nonsynonymous alterations in the Stage III/IV EPICA MM cohort, divided by methylation groups. Rows and columns represent genes and tumor lesions, respectively. Tumor Mutation Burden (TMB) and clinical features for each lesion are shown as tracks at the top. Frequencies of COSMIC mutational signatures are indicated as a bar plot at the bottom. The proportion of alterations among methylation groups for each gene is shown, with statistical significance indicated (p-value of Pearson’s chi-squared test statistic: *: p < 0.05, .: p < 0.1).

**Extended Data Figure 3.**
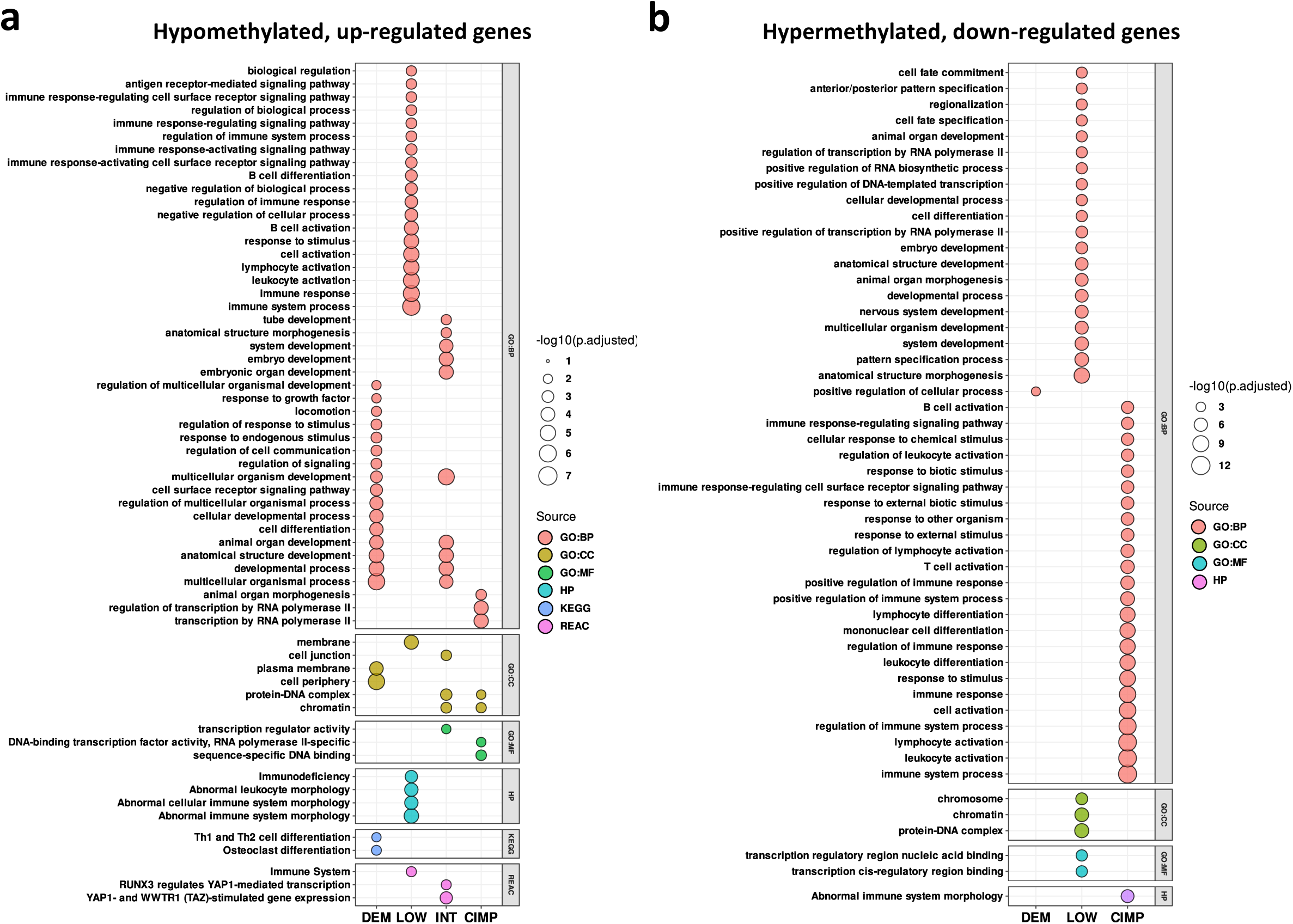
Promoter methylation vs. gene expression analysis. **a,b,** Over-representation analysis for association of promoter methylation vs. gene expression for selected gene sets. Dot plots showing the most statistically significantly enriched terms from over-representation analysis (ORA) of up-regulated genes whose promoters are hypomethylated (A) and down-regulated genes whose promoters are hypermethylated (B), comparing each methylation class vs all the others in EPICA cohort. Dots size represent the adjusted p-values and color codes represent collection of gene sets (GO:BP = Gene Ontology Biological Process; GO:CC = Gene Ontology Cellular Component; GO:MF = Gene Ontology Molecular Function; HP = The Human Phenotype Ontology; KEGG = Kyoto Encyclopedia of Genes and Genomes pathways; REAC = Reactome Pathway). Leukocyte fraction in B,C estimated according to: Thorsson, V. et al. The immune landscape of cancer. Immunity 48, 812-830 (2018).

**Extended Data Figure 4.**
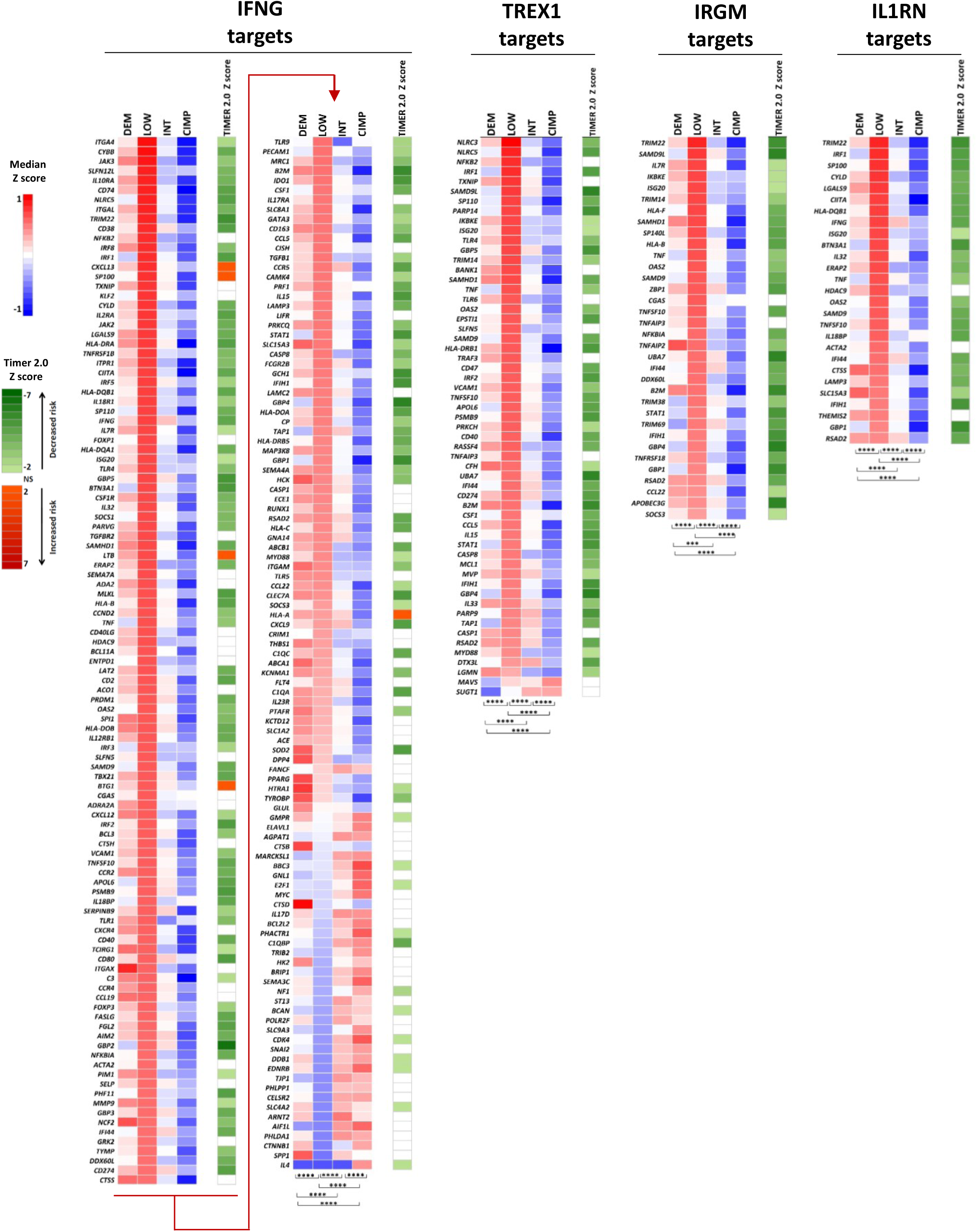
Expression and prognostic significance of the IFNG, TREX1, IRGM, and IL1RN target genes in EPICA methylation classes. Heatmaps of median Z score expression of IFNG, TREX1, IRGM, and IL1RN target genes in EPICA methylation classes. The column entitled «Timer 2.0 Z score» lists the Z score value indicating the prognostic significance of each gene in the TCGA MM cohort. Statistical analysis by Kruskal Wallis test followed by Dunn’s multiple comparison test. *: p<0.05; **: p<0.01, ***: p<0.001; ****: p<0.0001.

**Extended Data Figure 5.**
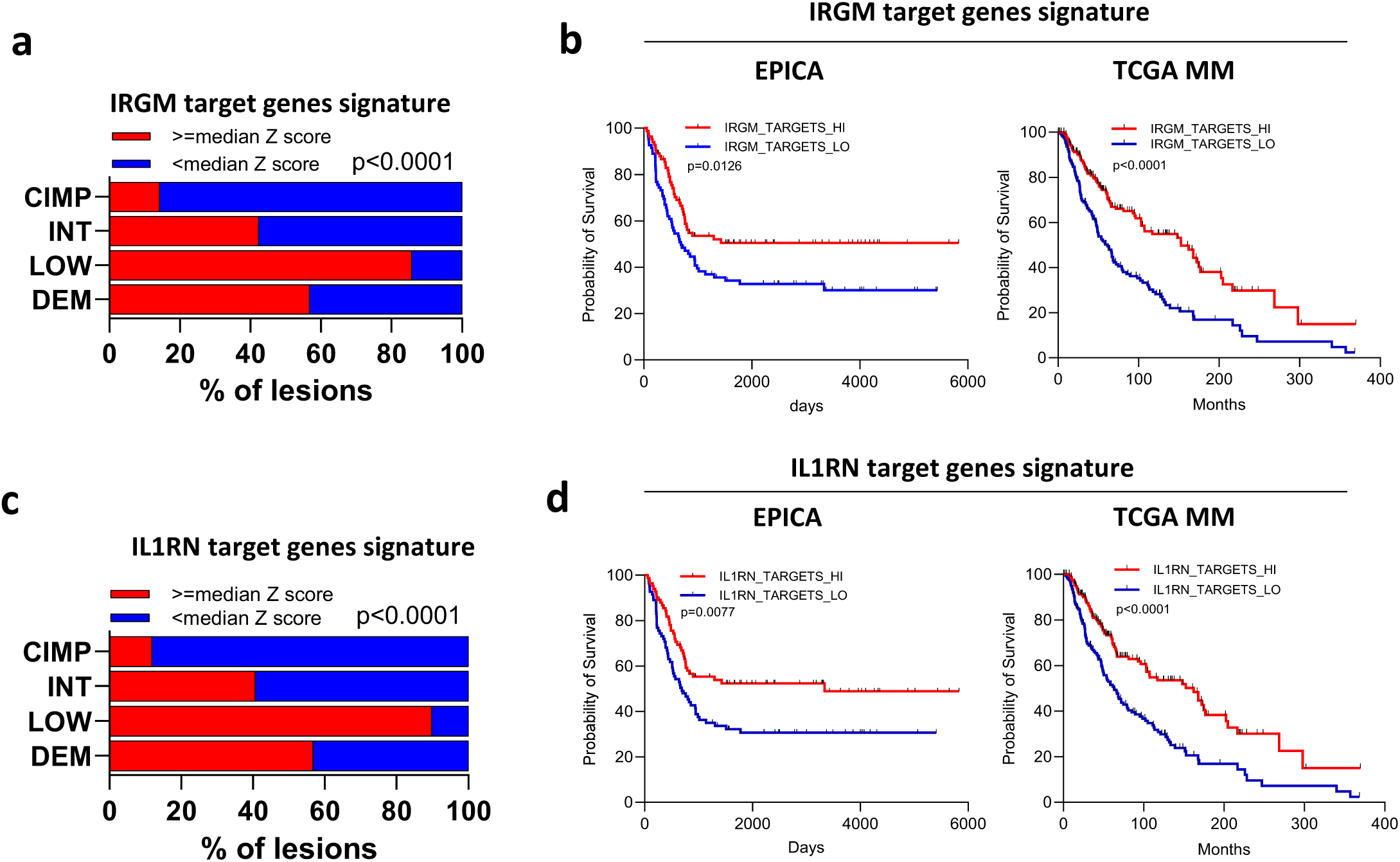
Top master molecules activated in CIMP MM classes are negative regulators of target genes with prognostic significance. **a,c,** Stacked bar plots showing, for each methylation cluster in the EPICA cohort, the percentage of samples with expression of the IRGM (a) or IL1RN (c) target genes above or below the median z score value of each signature. **b**,**d**, Kaplan-Meier survival curves of patients in the EPICA cohort (left plot) or TCGA MM cohort (right plot) according to median Z score expression (from RNA-seq profiling) of IRGM (b) or IL1RN (d) target gene signatures. In b,d patients in both cohorts were grouped according to median UR target gene expression above (“HI”) or below (”LO”) the median Z score value of the signature. Statistical analysis in a, c by chi square; in b, d by log rank test.

**Extended Data Figure 6.**
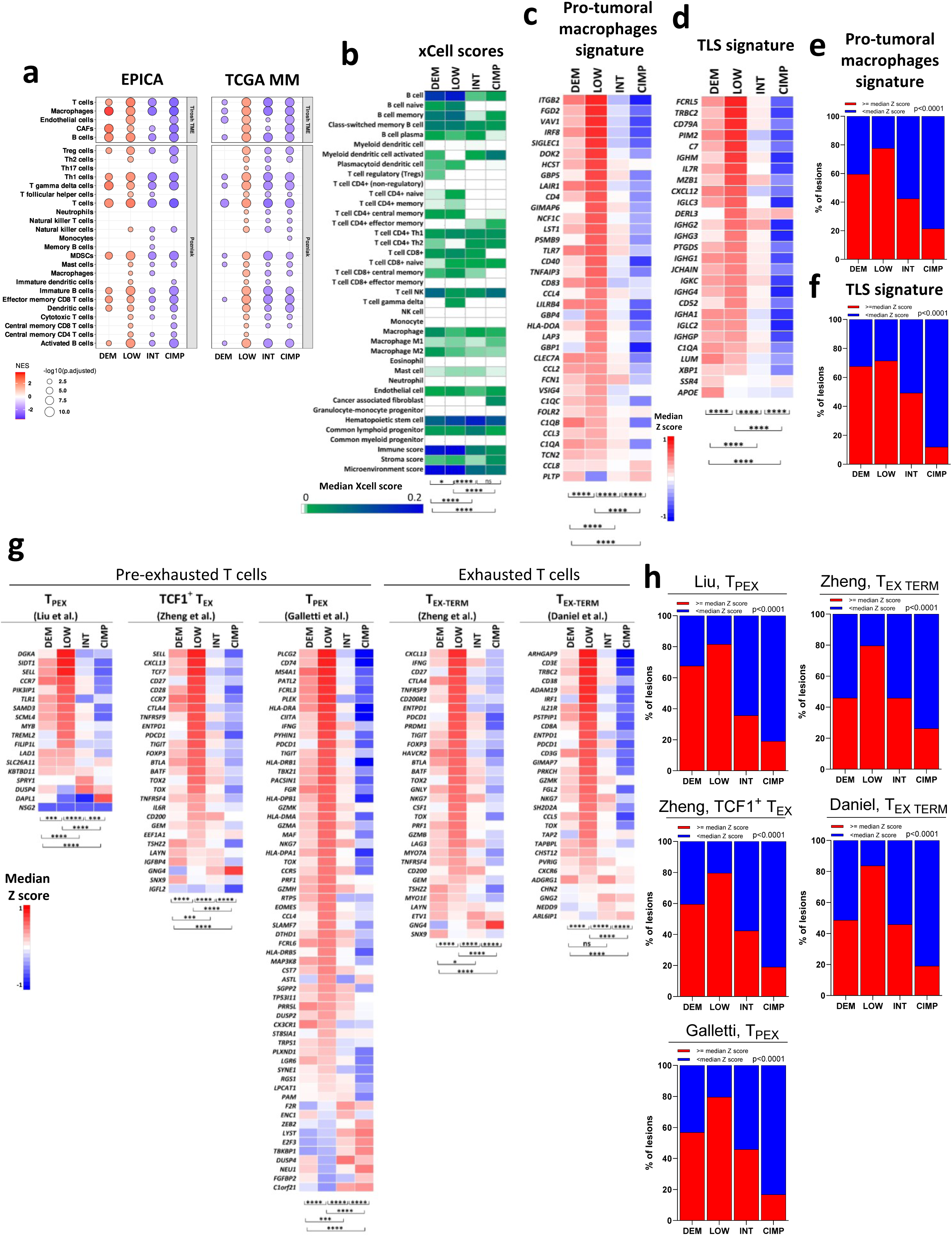
Assessment of immune-related gene signatures in the EPICA methylation-defined clusters. **a,** Dot plot of normalized enrichment scores (NES) of two melanoma microenvironment signatures^36–37^ in the four methylation subsets of the EPICA and TCGA metastatic melanoma cohorts. Dot size represents the adjusted p-values and scale colors represent the NES. **b,** Heatmap of median xCell algorithm scores in DEM, LOW INT and CIMP classes. **c,d,** Heatmaps of expression median z-scores in the four methylation subsets computed for genes belonging to the pro-tumoral macrophage signature^39^ (c) and TLS gene signature^40^ (d). **e,f,** Stacked bar plots showing for each methylation cluster the percentage of samples with expression of the pro-tumoral macrophages (e) or TLS (f) signatures above or below the median z score value of each signature. **g,** Heatmap of expression median z-scores in the four methylation subsets computed for genes belonging to three T_PEX_ signatures^41–43^ and to two T_EX_ signatures^43–44^. **h,** Stacked bar plots showing for each methylation cluster the percentage of samples with expression of the T_PEX_ and T_EX_ gene signatures above or below the median z score value of each signature. Statistical analysis by Kruskall Wallis test followed by Dunn’s multiple comparison test in b,c,d and g and by Chi-square in e,f,h.*: p<0.05; **: p<0.01, ***: p<0.001; ****: p<0.0001

**Extended Data Figure 7.**
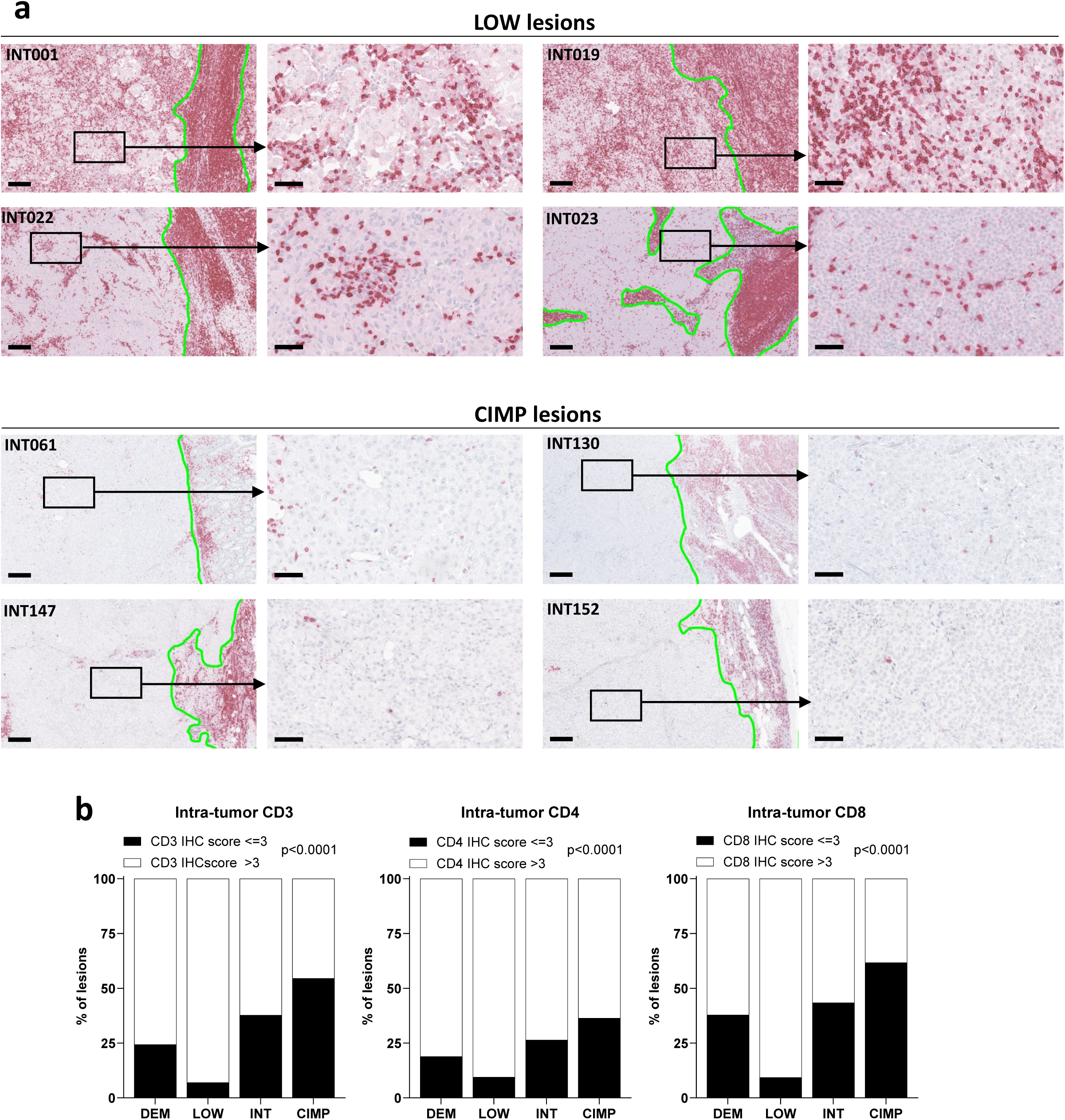
LOW lesions are enriched for intra-tumor T cells compared to CIMP lesions. **a**, representative images of four LOW lesions (TOP panels) and of four CIMP lesions (bottom panels) stained for CD3. For each lesion a higher magnification panel shows an intra-tumor area. The green line identifies the intra-tumor (left side of each image) and the extra-tumor (right side of each image) areas of each lesion. Scalebar: 200 μ (lower magnification image) and 50 μ (higher magnification image). **b**, Fraction of lesions in each methylation class having an intra-tumor CD3, CD4 and CD8 IHC score <=3 or >3. Statistical analysis by Chi-square.

**Extended Data Figure 8.**
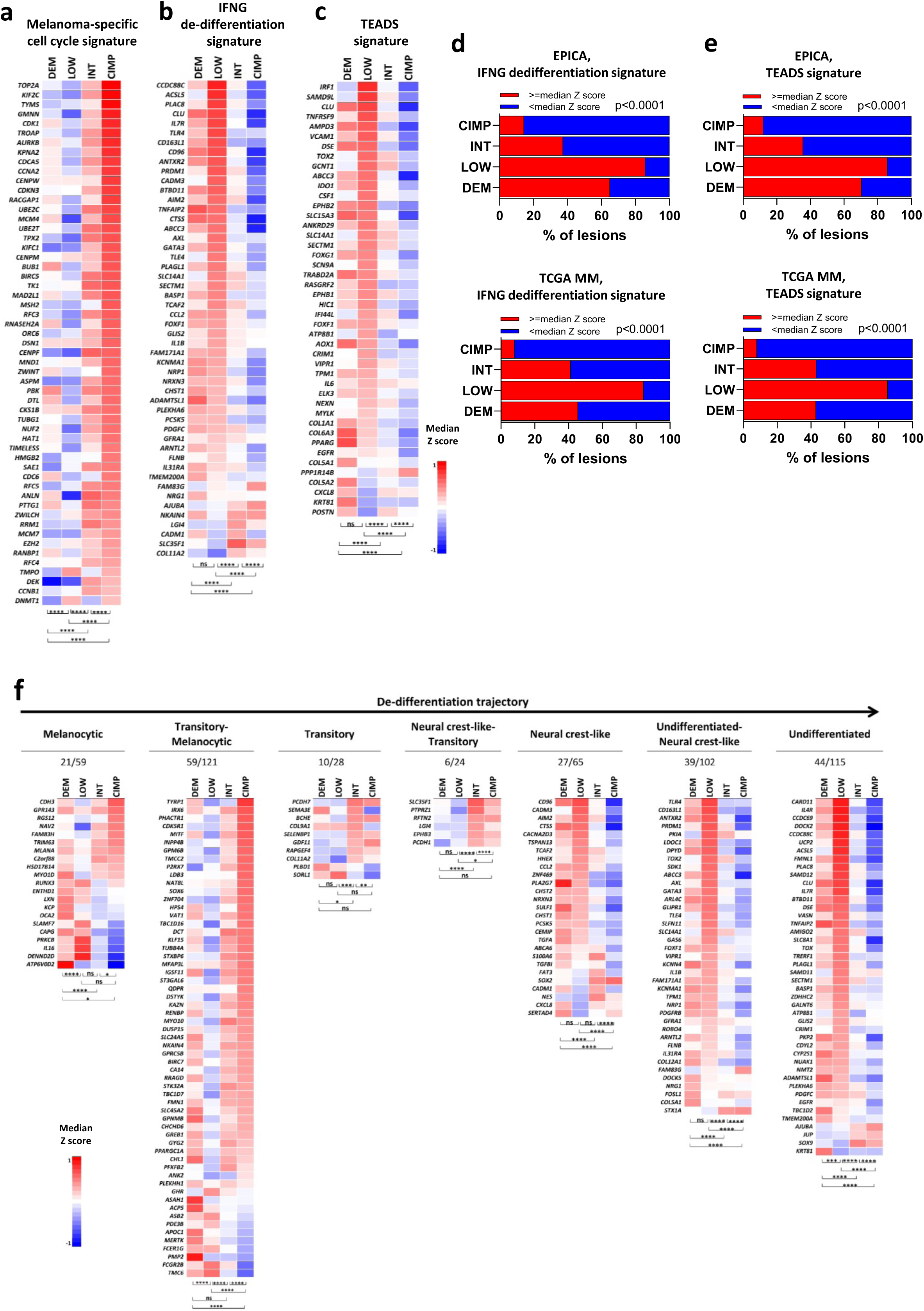
Expression of melanoma differentiation signatures in EPICA methylation subsets. **a,b,c,** Heatmaps of median Z score expression for genes in (a) of the melanoma-specific cell cycle signature^36^, in (b) of 50/111 genes in the IFNG de-differentiation signature^48^ and in (c) of 44/107 genes in the TEADS signature^49^ that discriminate the methylation defined classes of the EPICA cohort. **d,e,** Stacked bar plots showing for each methylation cluster in the EPICA cohort (Top graph) and TCGA metastatic melanoma cohort (bottom graph) the percentage of samples with expression of the IFNG de-differentiation (d) and TEADS signatures (e) above or below the median z score value of the signature. **f,** Heatmaps showing median z score values from RNA-seq profiling for selected genes in the seven sub-signatures describing melanoma differentiation according to Tsoi et al.^50^. Number of genes that discriminate methylation groups/total number of genes in each subsignature is shown below the sub-signature name. Statistical analysis in a, b, c, f: Kruskal Wallis test followed by Dunn’s multiple comparison test; in d,e by Chi-square. *: p<0.05, **: p<0.01; ***: p<0.001, ****: p<0.0001.

**Extended Data Figure 9.**
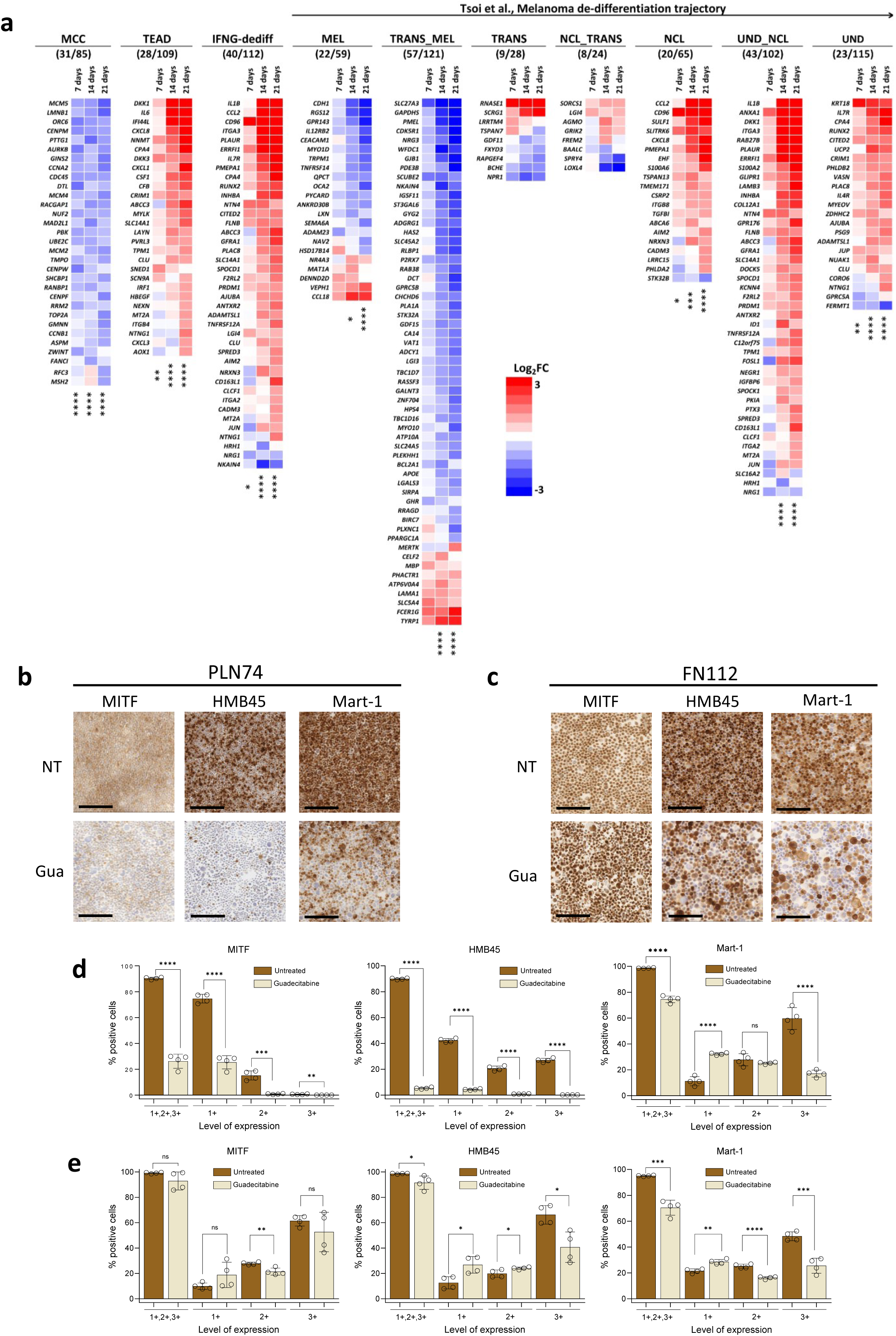
Guadecitabine treatment promotes melanoma de-differentiation. **a,** Heatmaps showing modulation of genes in the MCC^36^, TEADS^49^, IFNG-dediff^48^ and Tsoi et al.^50^ melanoma differentiation signatures in cell line PLN74 at day 7, 14 and 21 of treatment. For each gene signature only genes showing a Log_2_FC>|1| are shown. The number of these genes/the total number of genes in each signature is shown under the signature name. Statistical analysis by Mann Whitney test. **b,c.** Modulation of MITF, HMB45 and Mart-1 protein expression in two differentiated melanoma cell lines by guadecitabine. Images from cytospins of cell lines PLN74 (b) and FN112 (c) cultured or not with guadecitabine for 21 days and then stained with mAbs to MITF, HMB45 (PMEL/GP100) and MART-1 melanoma markers. Scale-bar = 200μm. **d,e,** Quantitative digital pathology analysis by QuPath software of MITF, HMB45 and Mart-1 expression in cell lines PLN74 (d) and FN112 (e) cultured or not with guadecitabine for 21 days. Results expressed as % positive cells at any level of expression (1+,2+,3+) or at levels 1+, 2+ or 3+. Statistical analysis in d,e by Student T test. *: p<0.05; **:p<0.01, ***:p<0.001; ****p<0.0001.

**Extended Data Figure 10.**
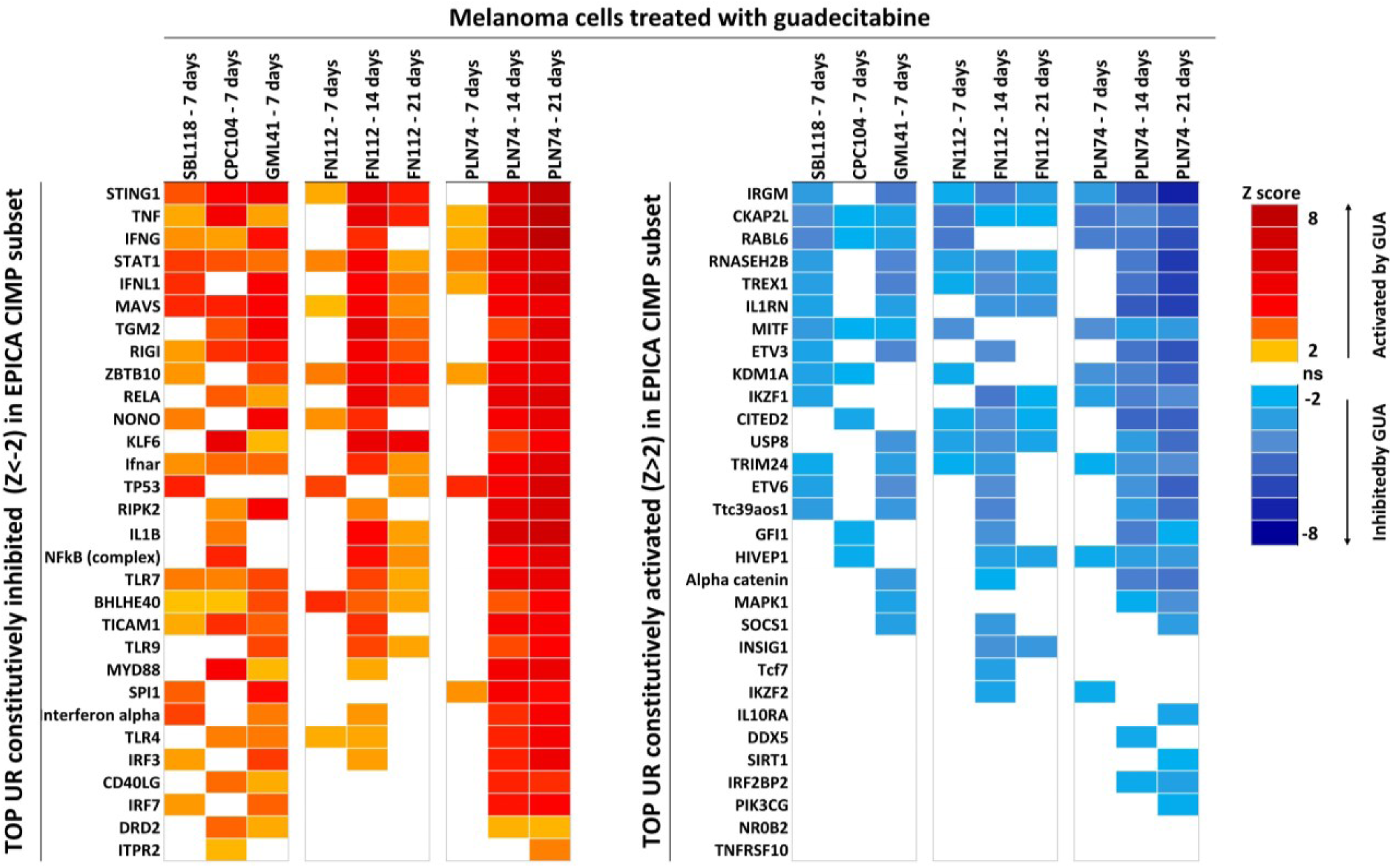
Guadecitabine treatment of differentiated melanoma lines reverses the activation state of UR identified in CIMP lesions. TOP UR constitutively inhibited in CIMP tumors (left) or constitutively activated in CIMP tumors (right) were selected and tested by IPA UR analysis for change in their activation state in melanoma cell lines treated at 7, 14 or 21 days with guadecitabine. The heatmaps show the significantly activated (orange to red) and significantly inhibited (light blue to blue) UR upon guadecitabine treatment based on z score values.

**Supplementary Figure 1.**
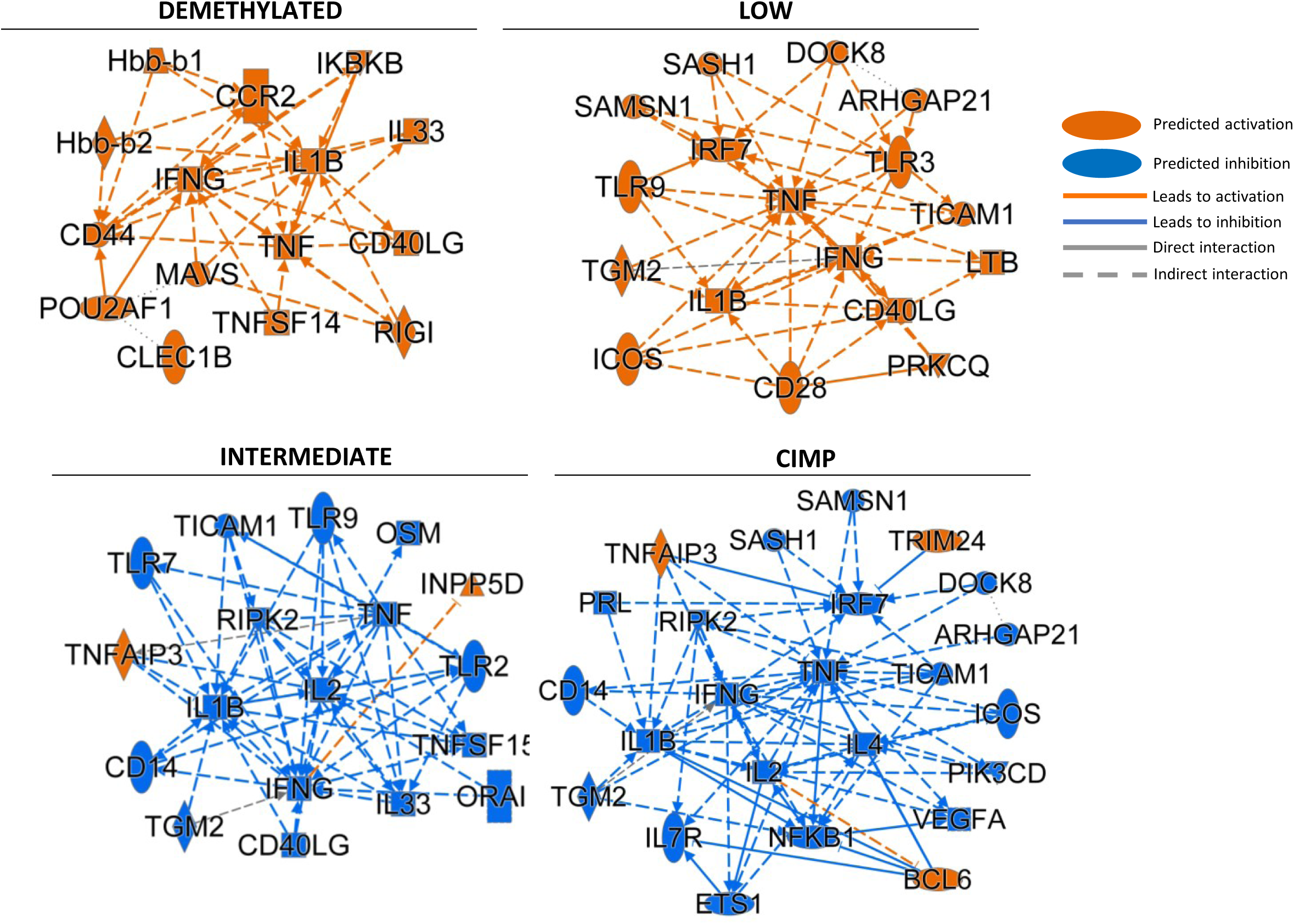
IPA core analysis for differentially expressed genes in the DEM, LOW, INT and CIMP EPICA classes. Graphical summary showing the most significant master molecules, their connections and relationships emerging from IPA core analysis for differentially expressed genes in the EPICA methylation classes. The orange and blue nodes mean “predicted activation” or “predicted inhibition”, respectively, of the indicated molecules; orange and blue arrows mean “leads to activation” or “leads to inhibition”, respectively; continuous and dashed lines mean direct or indirect interaction between main nodes, respectively; dotted line means inferred relationship between main nodes.

**Supplementary Figure 2.**
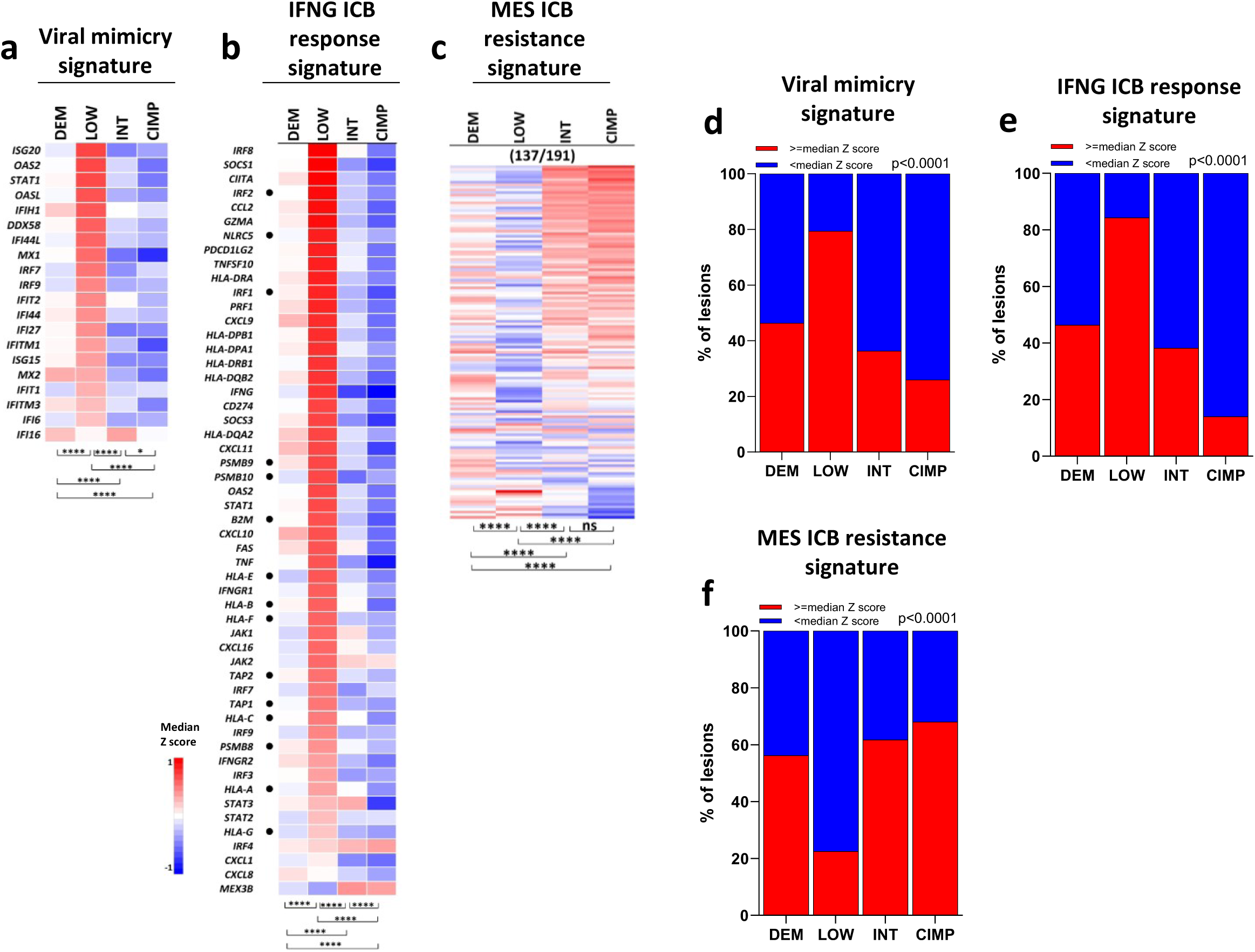
Expression of ICB predictive signatures in methylation-defined classes of the the TCGA MM cohort. **a,b,c,** Heatmaps of median Z score expression of genes in the viral mimicry (a), IFN ICB response (b) and MES resistance (c) signatures in methylation defined classes of the TCGA MM cohort. Genes identified by dots in panel B represent a core HLA Class I APM signature. **d**,**e**,**f**, Stacked bar plots showing for each methylation cluster in the TCGA MM cohort the percentage of samples with expression above or below the median z score value of each signature for the viral mimicry (d), the IFNG ICB response (e), and the MES resistance (f) signatures. Statistical analysis in a-c by Kruskal Wallis test followed by Dunn’s multiple comparison test; in d-f by Chi-square. *: p<0.05; **: p<0.01, ***: p<0.001; ****: p<0.0001.

**Supplementary Figure 3.**
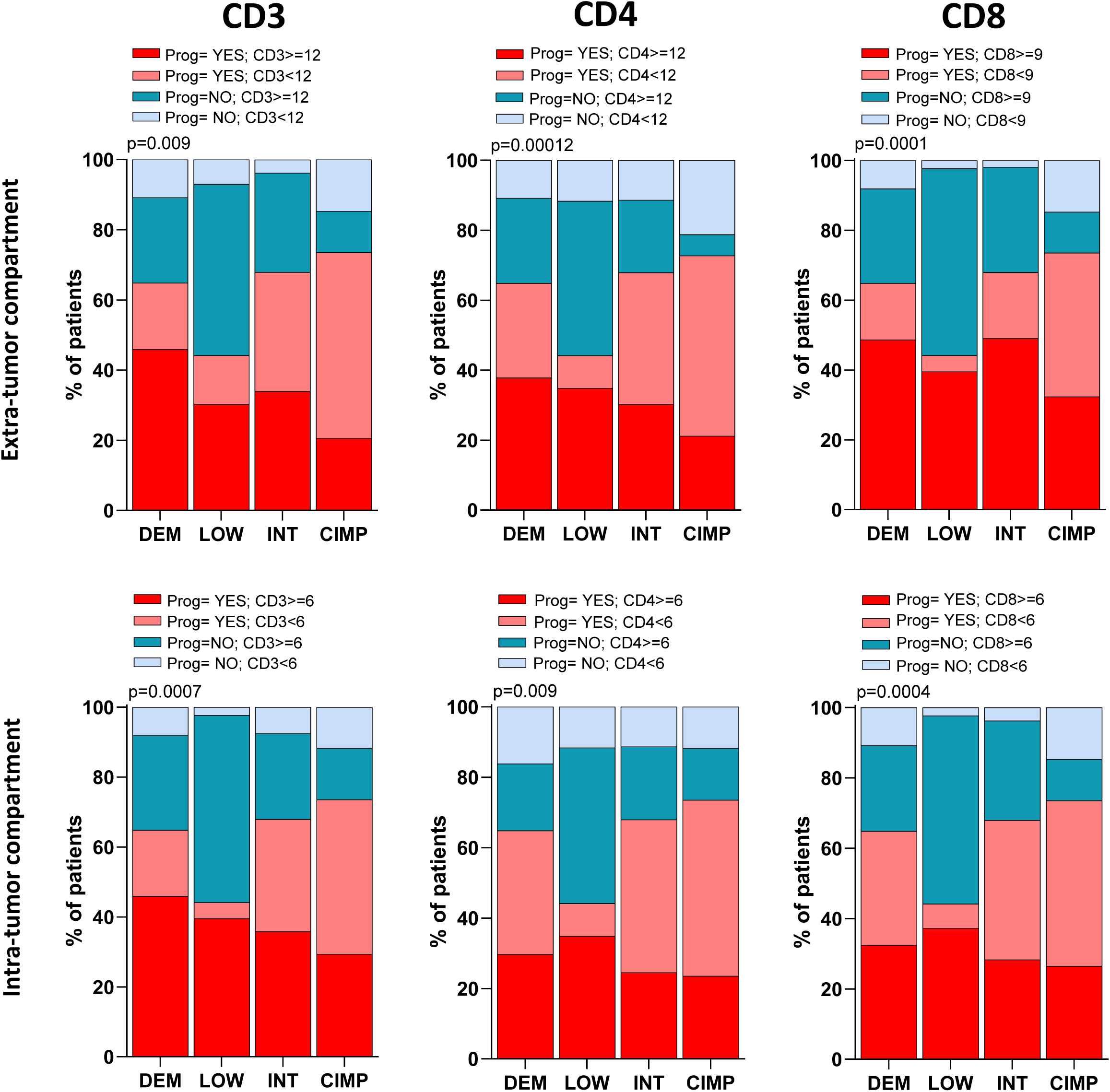
Association of methylation classes and tumor immune contexture with subsequent stage progression in EPICA cohort. Stacked bar plots of the percentages of EPICA patients, in each methylation class, grouped according to level of expression (above or below the median IHC score) of CD3, CD4 and CD8 in extra- or intra-tumor compartments in the initial investigated lesion and according to subsequent AJCC stage progression (Prog =YES, light red or dark red bars) or to lack of subsequent stage progression (Prog=NO, light blue or dark blue bars). Statistical analysis by Chi-square.

**Supplementary Figure 4.**
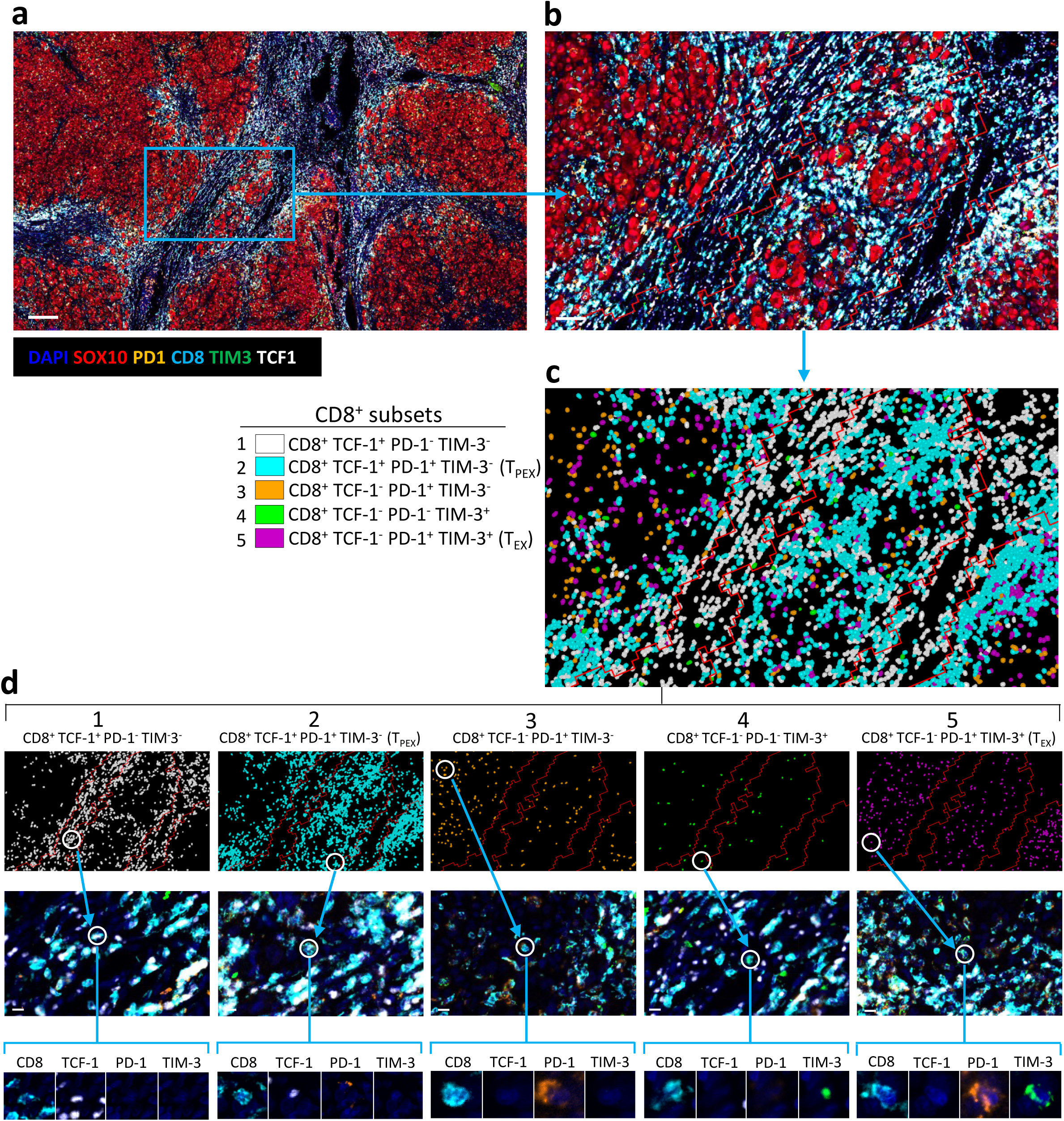
Analysis strategy for identification of CD8^+^ subsets characterized by differential expression of TCF-1, PD-1 and TIM-3 in EPICA melanoma lesions by multiple immunofluorescence (mIF). **a,** mIF image of a metastatic melanoma lesion (scale bar:200μ). **b,** A higher magnification area (scale bar: 100 μ) of the lesion in a. Red lines: tissue segmentation to discriminate tumor from stroma areas. **c,** Identity and position in the same area as in panel b, of CD8^+^ cells expressing the 5 indicated phenotypes. **d,** Upper panels: position of each of the identified CD8^+^ phenotypes in the same area as in c; middle panels: original mIF images centered on a higher magnification area containing representative examples of each CD8^+^ phenotype (scale bar: 10 μ); bottom panels: single channel mIF images showing expression of CD8, TCF-1, PD-1, TIM-3 in a single cell representative of each of the five identified subsets. Visualization of tumor cells was omitted in panel c and d.

**Supplementary Figure 5.**
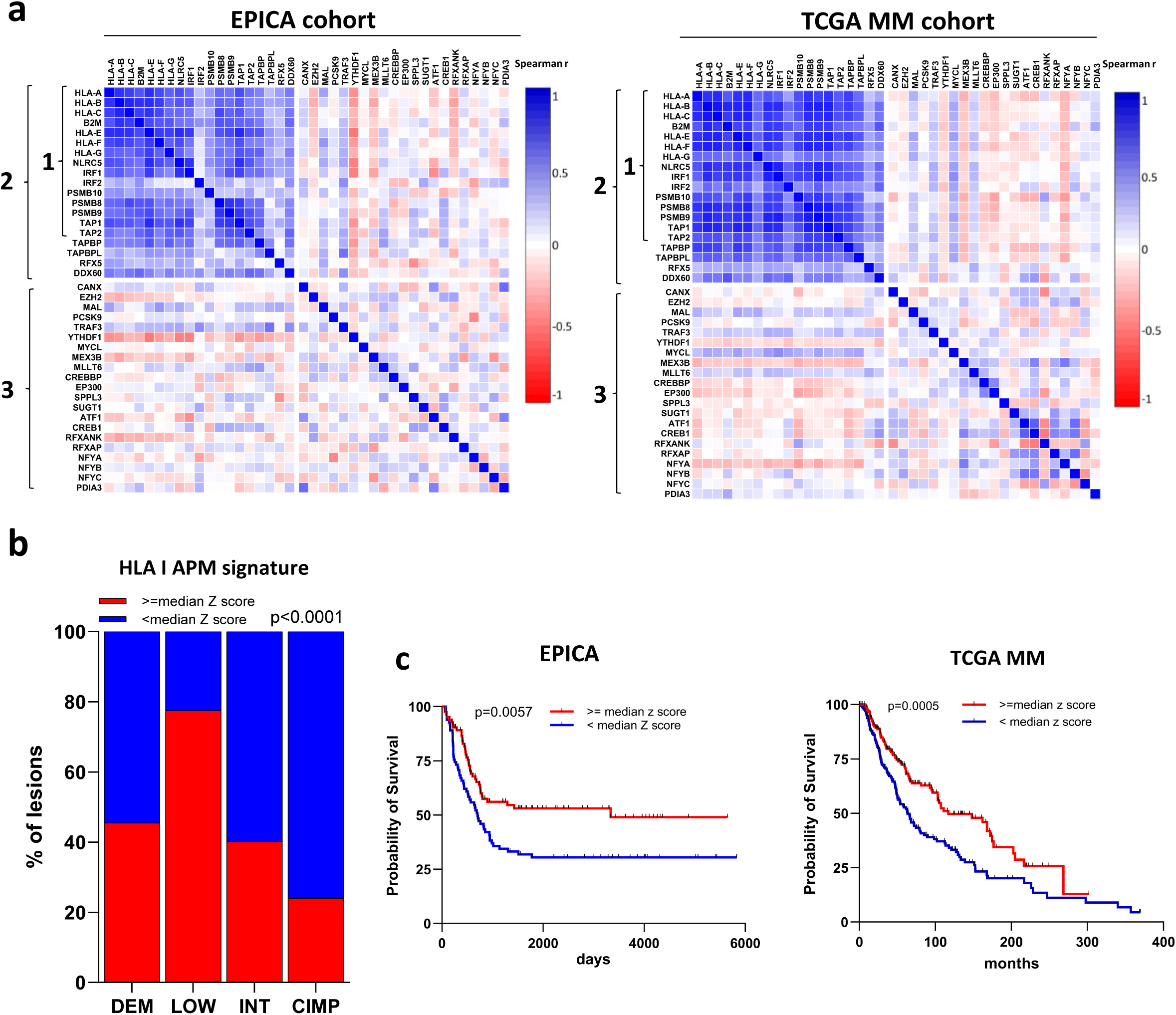
Correlation analysis in EPICA and TCGA MM cohorts of genes in the HLA Class I APM pathway and clinical significance of the HLA Class I APM signature. **a**, Spearman correlation analysis in EPICA and TCGA MM cohorts of the expression levels of genes in the core HLA class I APM signature (labelled “1”), in the extended HLA Class I APM signature (labelled “2”) and of genes involved in positive and negative regulation of HLA Class I expression (labelled “3”). **b,** Stacked bar plots showing for each methylation cluster in the EPICA cohort the percentage of samples with expression above or below the median z score value of the HLA Class I APM signature. **c,** Kaplan-Meier survival curves of patients in the EPICA cohort and in the TCGA MM cohort according to median z score value of the extended HLA Class I APM signature 2. Statistical analysis in a by spearman correlation, in b by Chi-square, in c by log-rank test.

**Supplementary Figure 6.**
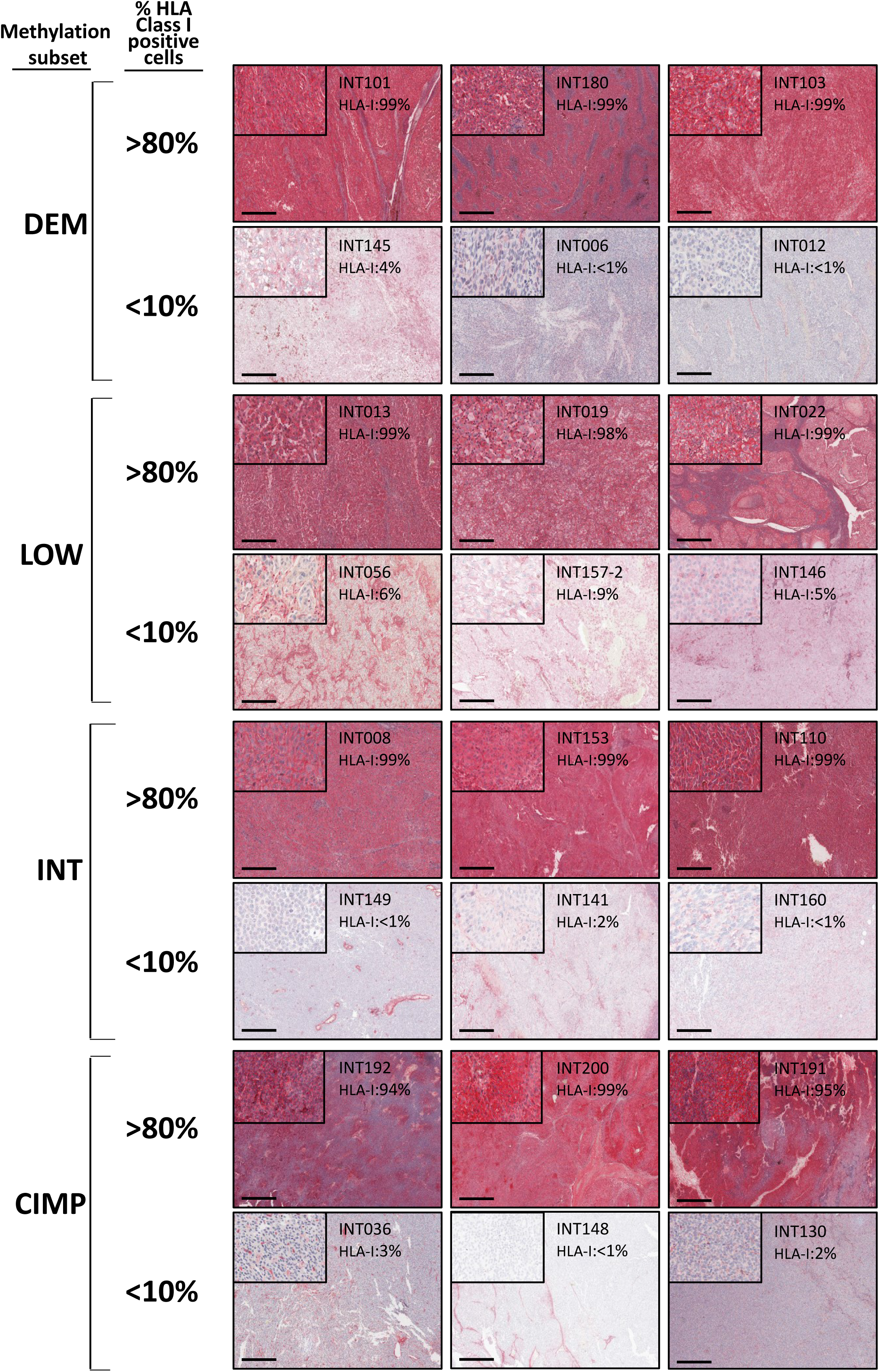
Expression of HLA Class I antigens on tumor cells in representative lesions belonging to the DEM, LOW, INT and CIMP classes of the EPICA cohort. Six representative lesions from each methylation class, stained in IHC for HLA Class I antigens and evaluated by quantitative digital pathology analysis are shown. For each methylation class three lesions with high expression of HLA Class I on tumor cells and three lesions with strong HLA Class I downmodulation are shown. For each lesion the inset shows a higher magnification of a tumor area. Scale bar: 500μ.

**Supplementary Figure 7.**
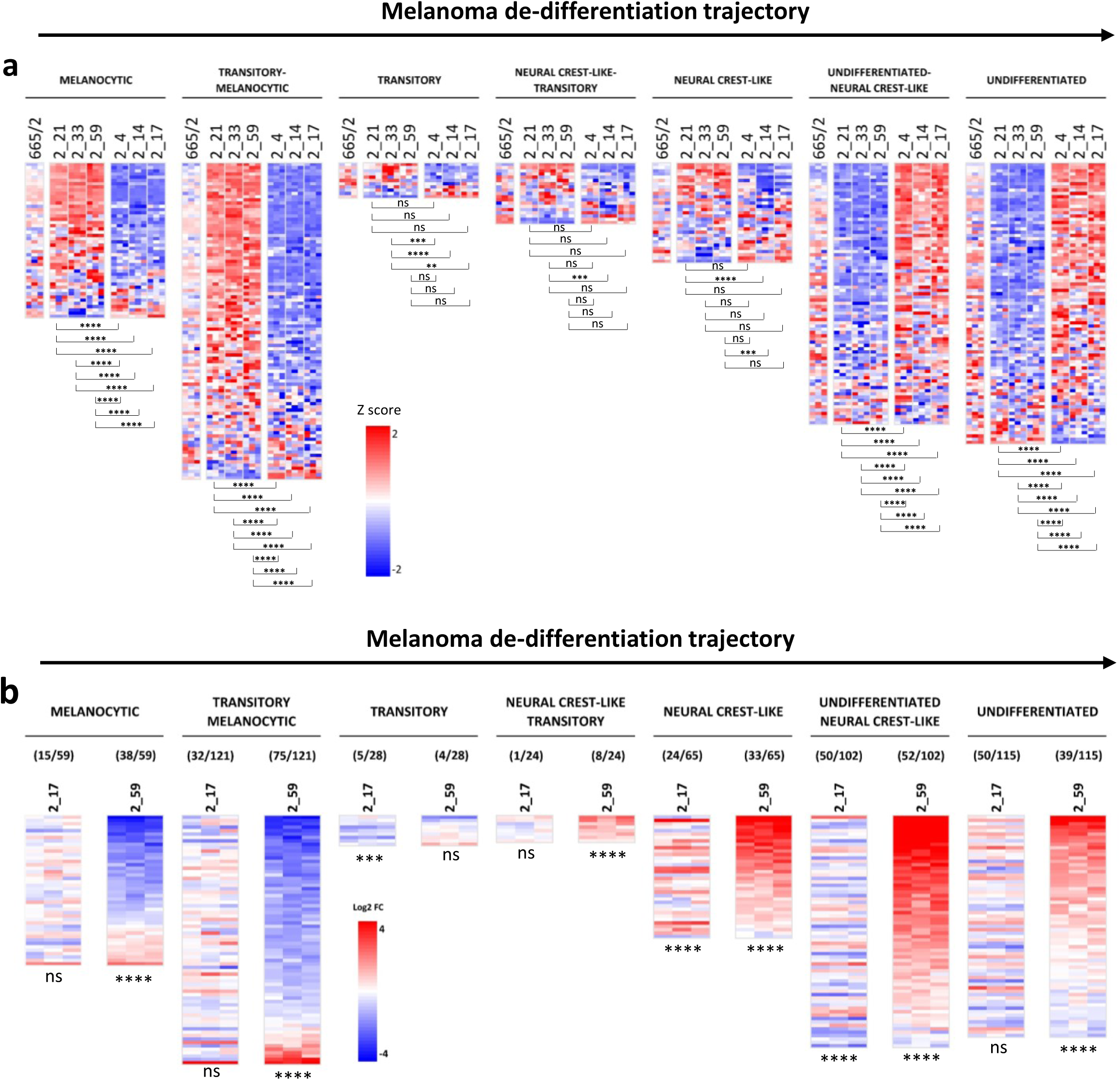
Guadecitabine promotes de-differentiation of the differentiated melanoma clone 2_59. **a,** Heatmaps showing expression of genes in the seven melanoma differentiation sub-signatures (Tsoi et al.^50^) in the parental line 665/2 and in six clones (2_21_2_33, 2_59, 2_4, 2_14, 2_17) generated from the parental line 665/2. Two groups of clones can be identified. For each sub-signature, only genes discriminating clones 2_21, 2_33 and 2_59 from clones 2_4, 2_14 and 2-17 are shown. **B,** Modulation of genes in the seven melanoma differentiation sub-signatures (Tsoi et al.^50^) by treatment with guadecitabine in melanoma clones 2_17 and 2_59. For each sub-signature, only genes showing significant modulation by guadecitabine in each of the two clones are shown. The number of these genes/the total number of genes in each sub-signature is shown under the signature name. Statistical analysis in a, b by two way ANOVA followed by Tukey’s multiple comparison test. *: p<0.05; **:p<0.01; ***:p<0.001; ****:p<0.0001.

