## Supplementary Methods for "Integrated multi-omics profiling reveals the role of the DNA methylation landscape in shaping biological heterogeneity and clinical behaviour of metastatic melanoma"

[1]Department of Experimental Oncology, Fondazione IRCCS Istituto Nazionale dei Tumori, Milan, Italy; [2]Biogem Institute of Molecular Biology and Genetics, Ariano Irpino, Italy; [3]Department of Electrical Engineering and Information Technologies, University of Naples Federico II, Naples, Italy; [4]Sylvester Comprehensive Cancer Center, and Department of Public Health Sciences, Miller School of Medicine, University of Miami, Miami, FL, USA; [5]University of Siena, Italy, and Center for Immuno-Oncology, Department of Oncology, University Hospital of Siena, Siena, Italy; [6]NIBIT Foundation Onlus, Italy; [7]Department of Advanced Diagnostics, Fondazione IRCCS Istituto Nazionale dei Tumori, Milan, Italy; [8]Section of Anatomic Pathology, Department of Health Sciences, University of Florence, Florence, Italy; [9]Department of Oral and Maxillofacial Surgery, New York University - College of Dentistry, New York, USA; [10]Department of Surgical Oncology, Fondazione IRCCS Istituto Nazionale dei Tumori, Milan, Italy.

[11]These authors contributed equally.

[§]These authors jointly supervised the work.

**Patient eligibility and lesion quality control.** Patients in the EPICA cohort (n=165) had diagnosis of cutaneous melanoma and underwent surgery for resection of metastatic lesions at Fondazione IRCCS Istituto Nazionale dei Tumori, Milan, between September 2002 and December 2017. Relevant demographic and clinicopathological data are listed in patient centric form in Supplementary Table 1A and in lesion centric form in Supplementary Table 1B**.** For all patients, survival was calculated from the date of surgery for resection of the investigated lesion to death, or to last follow-up. Age was calculated from date of birth to date of surgery for resection of the investigated lesion. For seventeen patients, multiple metachronous or synchronous lesions were investigated (range 2 to 6 lesions/patient, Supplementary Table 1A). No systemic treatments before surgery for resection of the first investigated lesion were received by 162/165 patients (98.2%, Supplementary Table 1A). Treatments after surgery for resection of the investigated lesions are summarized in Supplementary Table 1A and listed in patient centric form in Supplementary Table 1B. Hematoxilin and eosin (H&E) sections of all candidate lesions from all patients were reviewed by two expert pathologists and lesions to be investigated (n=191) were selected based on on appropriate biospecimen size, percent of tumor cells, extent of necrosis, amount of melanin pigment and tumor morphology. Mean tumor purity of the EPICA cohort was 83.1% (Supplementary Table 1B**)**. For the adjuvant cohort, patients' selection was based on the availability of adequate amount of formalin-fixed, paraffin-embedded (FFPE) tumor samples collected before ICB therapy and fully annotated medical history. For all patients, recurrence-free survival (RFS) was calculated from the date of first cycle of ICI therapy to the evidence of first recurrence (local or distant), or to date of the last follow-up or death. Age was calculated from date of birth to date of surgery for resection of the investigated lesion. No systemic treatments before surgical resection of the first investigated lesion were received by all patients. Relevant demographic and clinicopathological data of this cohort and treatments after surgical resection of the investigated lesions are listed in Supplementary Table 4.

**WES quality controls and data analysis**. Quality control of the WES data was performed using fastQC (v. 0.11.8) (https://www.bioinformatics.babraham.ac.uk/projects/fastqc/). Sequencing reads were aligned to the human reference genome (UCSC genome assembly GRCh37/hg19) using the Burrows-

Wheeler Aligner (1). Further processing with GATK (2) involved removing low mapping quality reads and realigning indels. Somatic single-nucleotide variants (SNVs) and indels were identified using the Sentieon Genomic Tool v. 201911 (3). In the absence of matched normal samples, both a virtual normal panel from the 1000 Genomes Project (4) and a pre-trained machine-learning model (5) were employed to ensure high-confidence calling of somatic SNVs and indels. Annotation of SNVs and indels was conducted using AnnoVar (6) and SnpEff (7), aggregating information from genomic and protein resources (GENECODE, UniProt, dbNSFP), cancer (COSMIC, ClinVar), and non-cancer variant databases (dbSNP, 1000 Genomes, Kaviar, Haplotype Reference Consortium, Exome Aggregation Consortium, NHLBI Exome Variant Server). Only variants affecting the protein sequence (missense, truncating, stop-loss, splicing variants, frameshift, and in-frame indels) were selected. Variants with a minor allele frequency ≥ 0.05 in non-cancer databases were excluded. Additionally, common sequencing artifacts, such as variants in very large genes (e.g., TTN and USH2A) and highly paralogous genes (e.g., mucins and keratins), were filtered out. The functional impact of missense SNVs and in-frame indels was predicted using PolyPhen-2 (8), SIFT (9), and PROVEAN (10), and variants predicted as damaging by at least two of these algorithms were classified as pathogenic. The nonsynonymous tumor mutational burden (TMB) was calculated as the number of nonsynonymous somatic mutations (single-nucleotide variants and small insertions/deletions) per megabase in coding regions. Frequencies of COSMIC mutational signatures v3.2 (11) were determined using the deconstructSigs R package (v. 1.8.0) (12).

**RRBS data analysis.** Only CpG sites outside regions annotated as “Open Sea” and with a minimum average of 10× coverage depth across all samples were selected for the analysis. Differential DNA methylated CpG sites between methylation subtypes were assessed using computeDiffTab.extended.site function in RnBeads package. Differential DNA methylated promoters between predefined groups of samples were assessed using the two-sided Welch t-test.

Only promoters with an adjusted p.value less than 0.10 (Benjamini & Hochberg method) were considered as differentially methylated. To derive methylation subtypes, the 1% most variable CpG sites were selected based on the interquartile difference at 0.9 and 0.1 percent. Consensus clustering was performed using the R package ConsensusClusterPlus (13). In details, 75% of the tumor samples were randomly subsampled without replacement (1000 fold) and partitioned into 4 major clusters using the K-means algorithm with Euclidian's distance as metric. The optimal number of clusters was determined by evaluating the relative change in area under the CDF curve for k = 2 to 8. For the adjuvant samples the classification of the cohort into the four methylation subtypes was performed using the k-nearest neighbors (kNN) algorithm from the R package class (version 7.3-22). The beta value matrix of pre-adjuvant samples was combined with that of metastatic melanoma samples from stages III and IV, retaining only the shared sites. The 191 metastatic melanoma samples were used as the training set to classify the pre-adjuvant samples, with the number of nearest neighbors (k) determined as the square root of the total number of samples in both the training and testing sets.

**Validation of methylation subtypes in TCGA cohort.** The DNA 450K methylation array and normalized gene expression data for TCGA skin cutaneous melanoma (SKCM) were retrieved from GDC Legacy Archive using TCGAbiolinks R package (14). Primary (n=104) and metastatic (n=368) SKCM samples were analyzed separately. DNA methylated probes annotated as multi-mapping, cross-reactive, "Open Sea" or mapping on X/Y chromosomes were filtered out. The 1% most variable CpG sites were selected based on the inter-percentile range between 90% and 10%. Unsupervised clustering of samples based on variable probes was performed using hierarchical clustering (complete linkage method and Euclidean distance).

**RNA-seq data analysis pipeline**. Raw data were normalized according to sample-specific GC-content differences as described in the EDAseq R package (v. 2.22.0) (15). Differential gene expression among groups of samples was estimated using the Mann-Withney-Wilcoxon test, and genes sorted according to its statistics were used for performing Gene Set Enrichment Analysis (GSEA) (16) of a curated collection of gene sets, as implemented in the clusterProfiler R package (v. 3.3.6) (17). The gene-set collections used for GSEA analysis included: (i) the GO Biological Process ontology from MSigDB (18) and (ii) a manually curated collection of signatures related to melanoma tumor microenvironment subpopulations. GO terms and signatures with an adjusted p-value less than 0.05 (Benjamini & Hochberg method) were considered significant. Additionally, single-sample gene set enrichment analysis of KEGG pathways from MSigDB (18) was performed using the mmw-GST function in the yaGST R package (19). KEGG pathways with an absolute logit2 normalized enrichment score (logit2-NES) greater than 0.58 and a nominal p-value less than 0.05 in at least 80% of samples were retained. Unsupervised clustering of EPICA cohort samples based on retrieved pathways was conducted using hierarchical clustering (complete linkage method and Euclidean distance). MIRACLE and IMPRES scores were computed using the MIRACLE R package as described in Turan et al (20) and calc impres R function (hhttps://github.com/Benjamin-Vincent-Lab/binfotron/) (21), respectively. For graphical purposes the scores were rescaled between 0 and 1. The differences within each group compared to all other groups were calculated using Student's T test, with p values adjusted to the benjamini-Hochberg procedure.

**Integrative analysis of RNA-seq and RRBS data.** Integration of gene expression and methylation data was performed used the SMITE R package (v. 1.16.0) (22) as previously described (23). For each methylation subtype, hypomethylated/upregulated and hypermethylated/downregulated genes were functionally characterized through a pathway enrichment analysis within the R package SMITE. Significant categories (p value < 0.01) were visualized as dot plot using R package ggplot2.

**Whole genome gene expression analysis by GeneChip Human Clariom S.** The quality of total RNA was first assessed using the RNA 6000 Assay RNA chips run on an Agilent Bioanalyzer 2100 (Agilent

Technologies, Palo Alto, CA). The total RNA (100 ng) was reverse transcribed using GeneChip 3'IVT Pico Reagent Kit (Affymetrix; Thermo Fisher Scientific, Inc.). As suggested by manufacturer's protocol, Poly-A RNA Controls were added to each sample, at a final concentration of *lys* 1:100,000, *phe* 1:50,000, *thr* 1:25,000 and *dap* 1:6,667. Each eukaryotic GeneChip probe array contains probe sets for several *B. subtilis* genes that are absent in the samples analyzed (*lys*, *phe*, *thr*, and *dap*). This Poly-A RNA Control Kit contains *in vitro* synthesized, polyadenylated transcripts for these *B. subtilis* genes that are pre-mixed at staggered concentrations to allow GeneChip probe array users to assess the overall success of the assay. The resulting cDNA was used as a template for obtaining cRNA; cRNA amplification is achieved by linear amplification using T7 in vitro transcription technology. The cRNA was then purified using Nucleic Acid Binding Beads (GeneChip 3'IVT Pico Reagent Kit, Affymetrix) and used as a template for reverse transcription followed by DNA polymerization to produce double-stranded DNA. During this step, dUTP was incorporated. The ds-cDNA was then fragmented using uracil-DNA glycosylase (UDG) and apurinic/apyrimidinic endonuclease 1 (APE 1) and labeled with DNA labeling reagent covalently linked to biotin using terminal deoxynucleotidyl transferase (TdT, GeneChip 3'IVT Pico Reagent Kit, Affymetrix). Hybridization of each fragmented and labeled target was performed using the GeneChip Hybridization, Wash and Stain Kit (Affymetrix; Thermo Fisher Scientific, Inc). It contains mix for target dilution, DMSO at a final concentration of 7% and pre-mixed biotin-labelled control oligo B2 and bioB, bioC, bioD and cre controls (Affymetrix cat #900299) at a final concentration of 50 pM, 1.5 pM, 5 pM, 25 pM and 100 pM, respectively. Targets were diluted in hybridization buffer at a concentration of 50 ng/ul and denatured at 99 °C for 5 minutes incubated at 45 °C for 5 minutes and centrifuged at maximum speed for 1 minute prior to introduction into the GeneChip cartridge. A single GeneChip Human Clariom S was then hybridized with each biotin-labeled target. Hybridizations were performed for 16 h at 45 °C in a rotisserie oven (60 RPM). GeneChip cartridges were washed and stained with GeneChip hybridization, Wash and Stain Kit in the Affymetrix Fluidics Station 450 following the S450_0007 standard protocol, including the following steps: (1) (wash) 10 cycles of 2 mixes/cycle with Wash Buffer A at 30 °C; (2) (wash) 6 cycles of 15 mixes/cycle with Wash Buffer B at 50 °C; (3) stain of the probe array for 5 min in SAPE solution at 35 °C; (4) (wash) 10 cycles of 4 mixes/cycle with Wash Buffer A at 30 °C; (5) stain of the probe array for 5 min in antibody solution at 35 °C; (6) stain of the probe array for 5 min in SAPE solution at 35 °C; (7) (final wash) 15 cycles of 4 mixes/cycle with Wash Buffer A at 35 °C; (8) fill the probe array with Array Holding buffer. GeneChip arrays were scanned using an Affymetrix GeneChip Scanner 3000 7G using default parameters. Affymetrix GeneChip Command Console software (AGCC) was used to acquire GeneChip images and generate .DAT and .CEL files, which were used for subsequent analysis with proprietary software. Gene expression data were analyzed by Transcriptomic Analysis Console (TAC) software (Applied Biosystems, Thermo Fisher Scientific). Analysis settings were as follows: gene level fold change >|1.2|, gene-level p value: <0.05; gene-level FDR: <0.05.

**Immunohistochemistry data analysis.** Images were acquired at ×20 with an Aperio Scanscope XT digital pathology slide scanner (Leica Biosystems). A Semi-quantitative analysis of the immune contexture was performed by an expert pathologist (MM) as reported (24, 25). Briefly, for intra- and extra-tumor compartments IHC data for each marker were rendered semi-quantitatively by adopting an IHC scoring system taking into account both staining marker extent (% positive cells) and intensity. The expression (E) was defined as follows: up to 25% neoplastic cells, 1+; 26–50%, 2+; 51–75%, 3+; 76–100%, 4+. The immunostaining intensity (I) was ranked as low (1+; fainter than internal controls), normal (2+; as faint as controls), or strong (3+; more intense than controls). E and I were combined into a single IHC score (S), calculated as E × I (24,25). For quantitative assessment of HLA Class I expression on tumor cells slides were scanned with Aperio Scanscope XT at 40X resolution and the whole tumor area was analyzed with QuPath v.0.5.0. Optimal thresholds were selected to identify negative, faint (1+), medium (2+) and strongly (3+) stained cells. Immunohistochemistry on melanoma cell lines was performed by setting up cytospins. Cytospins were prepared by resuspending the cell pellets in PBS using Shandon Cytospin 2 (Thermo Scientific) cytocentrifuge. Afterwards, samples were air dried and snap frozen. Slides thus obtained were briefly fixed in formalin (5 min), washed in PBS and subsequently stained with Leica Bond RX immunostainer (Leica Biosystems, Buffalo, IL) with a 5 minutes antigen retrieval with BOND Epitope Retrieval Solution 1. The following antibodies were used: Melan-A (Dako Agilent, clone A103), Melanosoma (Dako Agilent, clone HMB45) and MITF (Dako Agilent, clone D5). Slides were scanned with Aperio Scanscope XT at 40X resolution and the whole cytospin area was analyzed with QuPath v.0.5.0.

**mIF data analysis.** The whole slides were scanned at 20x using the Phenoimager HT scanner (Akoya Biosciences). Multispectral images were acquired using PhenoImager HT 2.0 software (Akoya Biosciences). Scans were unmixed with a melanoma-specific tissue library using Akoya's Inform software. Each image was analyzed using the open-source software QuPath v.0.5.0 (https://qupath.github.io/) (26). Images were then processed for tissue annotation, cell segmentation and cell phenotyping. Briefly, cell segmentation was based on Dapi nuclear staining using Stardist extension for QuPath v.0.4.0 (https://github.com/qupath/qupath-extension-stardist). To this end, a pixel classifier was created using machine learning algorithm within QuPath by exploiting S100/SOX10 signal to distinguish tumor epithelium and stroma; a phenotyping algorithm using Random Trees was developed for each marker using cytoplasmic or nuclear positive staining; the combined classifier was applied to each image. All steps of tumor-stroma distinction, cell segmentation, and phenotyping were supervised by a pathologist. Cell density (cells/$mm^2$) for each phenotype was calculated on tumor, stroma and total area.

**Upstream regulator analysis by IPA.** Ingenuity Pathway Analysis (IPA 8.5, www.ingenuity.com) was used to carry out Upstream Regulator (UR) analysis as described (27). UR analysis allows to identify upstream transcriptional regulators explaining the observed gene expression changes in the dataset. UR analysis returns results based on p-values and Z score statistics. P values indicate the likelihood of the association between a set of genes and related function, or the likelihood of the overlap between the genes in the dataset and those that are regulated by a predicted upstream regulator. The meaning of the Z score statistics is to infer the activation states ("increased" or "decreased") of the identified biological functions and of the predicted transcription factors. Only Z scores greater than 2 or smaller than -2 were considered significant.

**References.**
